# Head stabilization behavior and underlying circuit mechanisms in larval zebrafish

**DOI:** 10.1101/2025.11.12.688155

**Authors:** Takumi Sugioka, Tod R. Thiele, Herwig Baier, Masashi Tanimoto, Shin-ichi Higashijima

## Abstract

Head stabilization is essential for animal survival, enabling stable sensory input and effective motor coordination. Animals stabilize their heads in response to vestibular stimuli through the vestibulo-collic reflex (VCR). While the VCR has been characterized in tetrapod vertebrates, it remains unknown whether fish, which lack an anatomical neck, employ a comparable behavior. Here, we demonstrate that larval zebrafish exhibit VCR-like behaviors: they adjust their head orientation relative to the body by rostral body flexion during pitch tilts. The rostral body flexed ventrally during head-up posture, whereas it flexed dorsally during head-down posture. These flexions partially compensated for the head pitch changes, thereby contributing to head stabilization. We also identified the muscles and neural circuits responsible for these two types of body flexions. Both the dorsal and ventral flexions were mediated by the same vestibular nucleus, but neural signals were transmitted through distinct pathways, either involving or bypassing a class of reticulospinal neurons. The dorsal and ventral flexions were ultimately produced by specialized dorsal and ventral muscles in the rostral body, respectively. The neural circuits underlying these body flexions in fish share similarities with those underlying the mammalian VCR. Together, our results demonstrate that fish exhibit a VCR-like behavior through comparable circuit mechanisms, suggesting that the VCR is evolutionarily conserved across vertebrates.

## Introduction

Maintaining balance is important for most animals. In particular, head stabilization is critical for maintaining visual focus (e.g., gaze stabilization) and for coordinating movements^1,2^. The vestibular system plays a central role in controlling head orientation^3–5^. Head movements are detected by the vestibular organs in the inner ear, processed in the brainstem, and relayed to neck muscles to stabilize the head through the vestibulo-collic reflex (VCR)^6,7^. While the VCR has been characterized in tetrapod vertebrates^7–10^, it remains unknown whether fish also exhibit the VCR.

Fish and mammals differ in the structural connection between the head and body. In mammals, a neck exists between the skull and the pectoral (shoulder) girdle—for example, in humans, the scapula and clavicle^11^. This arrangement enables independent head movement relative to the body. By contrast, in fish, the skull is directly attached to the pectoral girdle^11,12^. Therefore, fish have been considered to lack an anatomical neck. Nevertheless, some fish species have been shown to possess cervical-like vertebrae and muscles homologous to mammalian neck muscles^13,14^. In addition, fish also exhibit “neck-like” movements, such as the rotation of their head relative to the body during feeding^15^. These findings raise the possibility that fish may exhibit a VCR-like behavior to stabilize the head in response to vestibular inputs. However, the associated behaviors and neural circuits remain entirely unknown.

To investigate whether fish exhibit a VCR-like behavior, we used larval zebrafish, an advantageous model organism for investigating the vestibular-related behaviors and neural mechanisms^16–18^. They have simple muscular and neural systems compared to adult fish, and their transparency facilitates the measurement and manipulation of neural activity^19,20^. Recent technical advances have substantially expanded our understanding of vestibular-related neural systems^21–26^. We previously reported that, in response to roll tilts, fish exhibit lateral body flexion that restores a dorsal-up posture^27,28^. Using a tiltable imaging system that enables Ca^2+^ imaging of tilted fish^23^, we also revealed the neural mechanisms underlying the lateral body flexion behavior^27^. Building on this work, we next investigated whether larval zebrafish exhibit VCR-like behaviors in response to pitch tilts. If so, we also aimed to identify the neural circuits underlying the behaviors.

Here, we investigated behaviors of the “neck” region in response to pitch tilts. We first showed that fish precisely controlled the head orientation relative to the body, depending on the pitch angle. During head-up posture, fish flexed the rostral body ventrally, whereas they flexed it dorsally during head-down posture. We next identified the neural circuits and muscles underlying these behaviors. Ventral and dorsal flexions were mediated by the same vestibular nucleus (tangential nucleus, TAN), but their neural signals were relayed through distinct pathways, either involving or bypassing a class of reticulospinal neurons (neurons in the nucleus of the medial longitudinal fasciculus, nMLF). Each flexion was produced by specialized muscles located ventrally (posterior hypaxial muscles; PHMs) or dorsally (supracarinalis anterior muscles; SCAs) in the rostral body. The behaviors and underlying circuits share similarities with the mammalian VCR. Together, our results show that fish also exhibit VCR-like behaviors and suggest that the VCR may be conserved across vertebrates.

## Results

### Larval zebrafish adjust the flexion around the “neck” region depending on the head pitch angle

To examine whether fish control head orientation relative to the body in response to pitch tilt stimuli, we first observed the behavior of freely moving fish. A 6-dpf larva was placed in a thin chamber, and its behavior was recorded from the lateral side under dark conditions (Figure 1A, left). Fish behavior alternated between swimming and non-swimming periods^29^ (the latter hereafter referred to as “pauses”). We noticed that the rostral body shape slightly changed during pauses depending on head pitch orientation. To quantify this, we defined two parameters: (1) the pitch angle, measured as the angle between the horizontal line and a line connecting the eye and the swim bladder (positive values indicate head-up posture, while negative values indicate head-down posture; Figure 1A, middle); and (2) the body bend angle, measured as the angle between lines fitted to the head and trunk contours (negative values indicate ventral bending; Figure 1A, right; see Methods for details). Figure 1B shows the raw images during pauses at different head pitch orientations (top), and rotated images, which enable comparison of the body shapes (bottom). When the fish was in a horizontal posture, the body slightly bent ventrally with the body bend angle being approximately −5° (Figure 1B, left). During head-up posture, the ventral bend became more pronounced and the body bend angle became around –10° (Figure 1B, middle). In contrast, during head-down posture, the body was nearly straight and the body bend angle became close to 0° (Figure 1B, right). For each pause, we calculated the mean pitch angle and mean body bend angle throughout the pause duration. In the population data, there was a strong correlation between these two angles across pauses (Figure 1C, *r* = −0.78; 340 pauses from 12 fish). In addition to pauses, we also analyzed the angles during swimming. As in pauses, a correlation was observed, with the mean body bend angle during swimming correlating with the pitch angle that was measured immediately before swimming (Figure S1A). These results suggest that fish finely adjust the rostral body in the dorso-ventral axis depending on head pitch angle.

**Figure 1.**
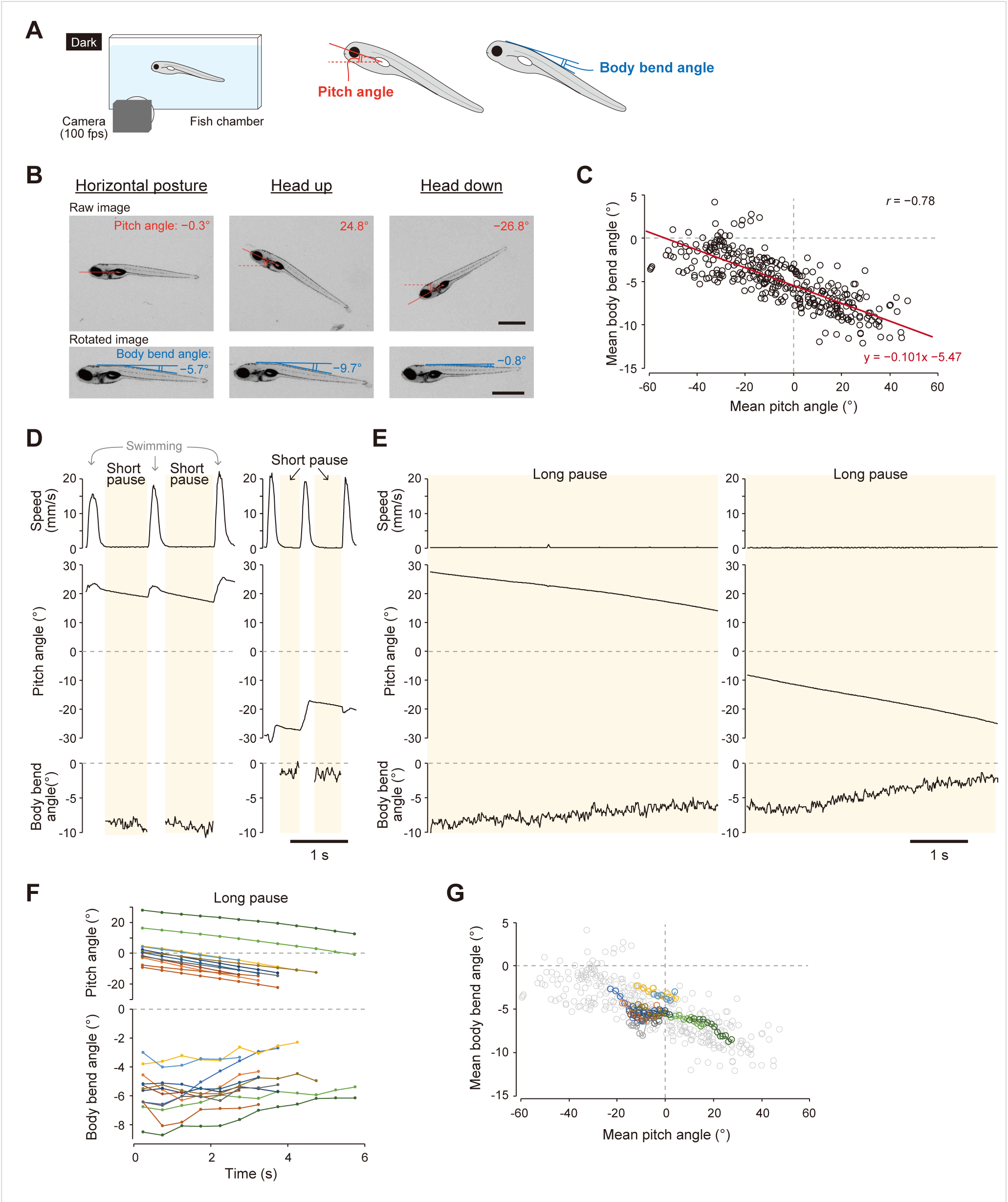
Body flexion in freely-moving fish. (A) Left: schematic of behavioral imaging. Middle: measurement of the pitch angle, defined as the angle between the horizontal line and a line connecting the center of the eye and the center of the swim bladder. Right: measurement of the body bend angle, defined as the angle between lines fitted to the head and trunk contours. (B) Snapshots of a fish with horizontal, head-up, and head-down postures during pauses. Top: raw images. Bottom: rotated images to facilitate comparison of the body shapes. Scale bars, 1 mm. (C) Body bend angles plotted against pitch angles during pauses. Both angles represent mean values calculated during each pause. Pauses longer than 0.2 seconds were analyzed (340 pauses from 12 fish). A regression line is shown in red. *r* represents Pearson correlation coefficient. (D) Example traces of speed, pitch angle, and body bend angle during short pauses with head-up (left) and head-down postures (right). Short pauses are indicated by yellow shading. Body bend angles were measured only during pauses. (E) Same as (D), but for long pauses that last over 3 seconds. (F) Traces of pitch angle and body bend angle during long pauses. Data points represent mean values calculated in 0.5-second bins (12 series from 5 fish). Each long pause series is shown in a different color. (G) Body bend angles plotted against pitch angles during long pauses. Data points represent mean values calculated every 0.5-second bin. The colors represent the same long pause series as in (F). The data are overlaid on the graph of short pauses shown in (C).

Fish are shown to passively rotate toward a head-down direction during pauses because the center of mass (where gravity acts) is located rostral to the center of volume (where buoyancy acts)^29^. Combined with our observations described above, this raises the possibility that the body bend angle increases as fish pitch down during a pause. To test this possibility, we analyzed the time course of the pitch and body bend angles throughout the pauses. The duration of pauses was, in most cases, less than 1 second (short pauses), but occasionally exceeded 3 seconds (long pauses) (Figure S1B). Figure 1D and 1E show speed, pitch angle, and body bend angle during short and long pauses, respectively. In both figures, the left panels correspond to head-up postures, and the right panels correspond to head-down postures. Consistent with a previous study by Ehrlich and Schoppik^29^, the pitch angle gradually decreased during pauses (Figure 1D and 1E, middle panels, yellow shading). The body bend angle did not increase during short pauses (Figure 1D, bottom). However, as expected, the angle showed a gradual increasing trend during long pauses (Figure 1E, bottom). Data from multiple long pauses further supported the hypothesis that the body bend angle gradually increased as fish pitched down, particularly in the late phase (> 2 s; Figure 1F). When the body bend angle was plotted against the pitch angle at each time point, the relationship closely matched that observed in the individual pauses (Figure 1G). The observation that the body bend angle did not noticeably increase during short pauses or the initial phase of long pauses suggests that it may be influenced by swimming activity immediately after movement cessation. Our freely moving behavioral analyses suggest that fish continuously adjust the rostral body shape in response to pitch angle.

### Head- or body-restrained fish exhibit dorsal and ventral flexions in response to pitch tilt stimuli

The modulation of the body bend angle depending on small pitch angle changes suggests that greater changes in head pitch angle would result in more pronounced changes in the body bend. To test this, we restrained either the head or the body of a fish and applied tilt stimuli. When the head was restrained, the tail was expected to shift dorsally or ventrally depending on the direction of the pitch tilt stimuli. Conversely, when the tail was restrained, the head was expected to shift accordingly.

We first restrained the head with agarose while allowing the body to move freely (Figure 2A). The fish was placed with their pitch angle approximately 0°, and was then tilted by 45° in either the head-up or head-down direction. During the initial horizontal posture, the body bend angle was negative, as in freely moving fish (Figure 2B, left; S1C; this value was slightly smaller, likely due to the embedding procedure). As expected, the body bend angle clearly changed in response to the pitch tilt stimuli (Figure 2B–2D). During head-up tilts, the trunk and tail shifted ventrally (Figure 2B, middle; 2C, top; 2D, left; Movie S1), whereas during head-down tilts, they shifted dorsally (Figure 2B, right; 2C, bottom; 2D, right; Movie S2). To quantify these postural changes, we focused on the changes in the body bend angle, and defined these behaviors as “ventral flexion” and “dorsal flexion,” respectively (Figure 2C, orange). Hereafter, “body bend angle” refers to the measured angle itself, while “flexion angle” denotes the relative change in the body bend angle from the horizontal posture (Figure 2D). Both ventral and dorsal flexions were consistently observed across multiple trials and different fish (Figure 2D, 2E).

**Figure 2.**
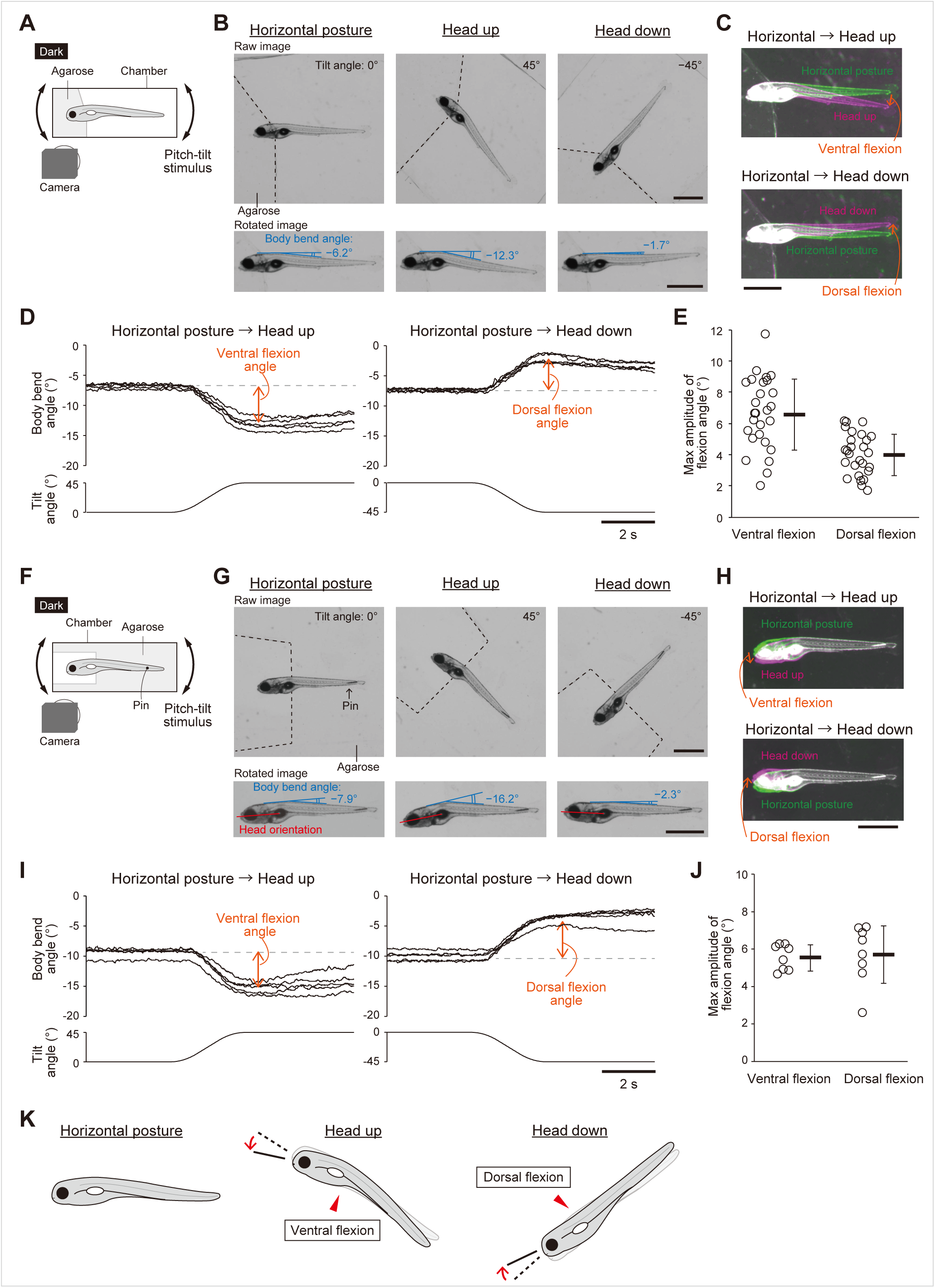
Body flexion in head- or body-embedded fish. (A–E) Body flexion in head-embedded fish (A) Schematic of behavioral imaging of head-embedded fish. (B) Snapshots of a head-embedded fish in horizontal, head-up, and head-down postures. Top: raw images. Black dashed lines indicate the edges of agarose. Bottom: rotated images. Image registration was performed using the head region to facilitate comparison of body shapes. (C) Composite images created from the rotated images of a fish in the horizontal (green) and tilted postures (magenta). (D) Example traces of body bend angles in response to tilt stimuli. Five trials for each tilt stimulus condition from a single fish are shown. Dotted lines indicate the baseline. The relative change in body bend angle from the baseline was defined as the ventral or dorsal flexion angle, as shown in orange. (E) Maximum amplitudes of ventral and dorsal flexion angles in naïve animals (27 fish). For each fish, the mean value of two to five trials is plotted. The thick line indicates the mean, and thin lines represent standard deviation. (F–J) Body flexion in body-embedded fish (F) Schematic of behavioral imaging of body-embedded fish. (G) Same as (B), but for a body-embedded fish. In the rotated images, image registration was performed using the tail region to facilitate comparison of head orientations. Red lines indicate head orientations. (H) Same as (C), but for a body-embedded fish. (I) Same as (D), but for a body-embedded fish. (J) Same as (E), but for body-embedded fish (8 fish). For each fish, the mean value of four to five trials is plotted. (K) Summary of behavioral experiments. Left: in a horizontal posture, the body slightly bends ventrally. Middle: during head-up tilts, fish flex the rostral body ventrally relative to the horizontal posture (ventral flexion), which contributes to bringing the head orientation closer to horizontal. The dashed line and transparent body indicate the head orientation and body shape that would occur without the ventral flexion. The head orientation change resulting from the ventral flexion is indicated by the solid line and arrow. Right: during head-down tilts, fish flex the rostral body dorsally relative to the horizontal posture (dorsal flexion), which also contributes to bringing the head orientation closer to horizontal. Scale bars, 1 mm.

We next restrained the body with agarose and a pin, allowing the head to move freely, and applied the same tilt stimuli (Figure 2F). Similar to head-embedded fish, the head shifted ventrally during head-up tilts and dorsally during head-down tilts (Figure 2G–2J). These flexions altered the head orientation relative to the body (Figure 2G, red; 2H), indicating that the head was tilted less than the applied tilt stimuli. Together, these head- or body-restrained experiments show that fish flex the rostral body ventrally or dorsally during head-up or head-down postures, respectively, and demonstrate that these body flexions partially compensate for the head pitch changes (Figure 2K).

### Specialized muscles on the dorsal and ventral sides of the “neck” region are activated during pitch tilts

Next, we explored muscles involved in the body flexion. It is reasonable to assume that the dorsal and ventral flexions recruit dorsal and ventral muscles near the flexion sites. These could be either trunk muscles that are typically recruited during swimming, or specialized muscle groups. If the latter is the case, a strong candidate muscle for the ventral flexion is the *smyhc2*-expressing posterior hypaxial muscles (PHMs)^27^, also known as the inferior obliquus muscles^30^ (Figur 3A). The *smyhc2* transgenic line also labels the supracarinalis anterior muscles (SCAs)^30^ (Figure 3A), which are located on the dorsal side of the rostral body and are well positioned to mediate the dorsal flexion. These data raise the possibility that *smyhc2*-expressing SCAs produce the dorsal flexion.

**Figure 3.**
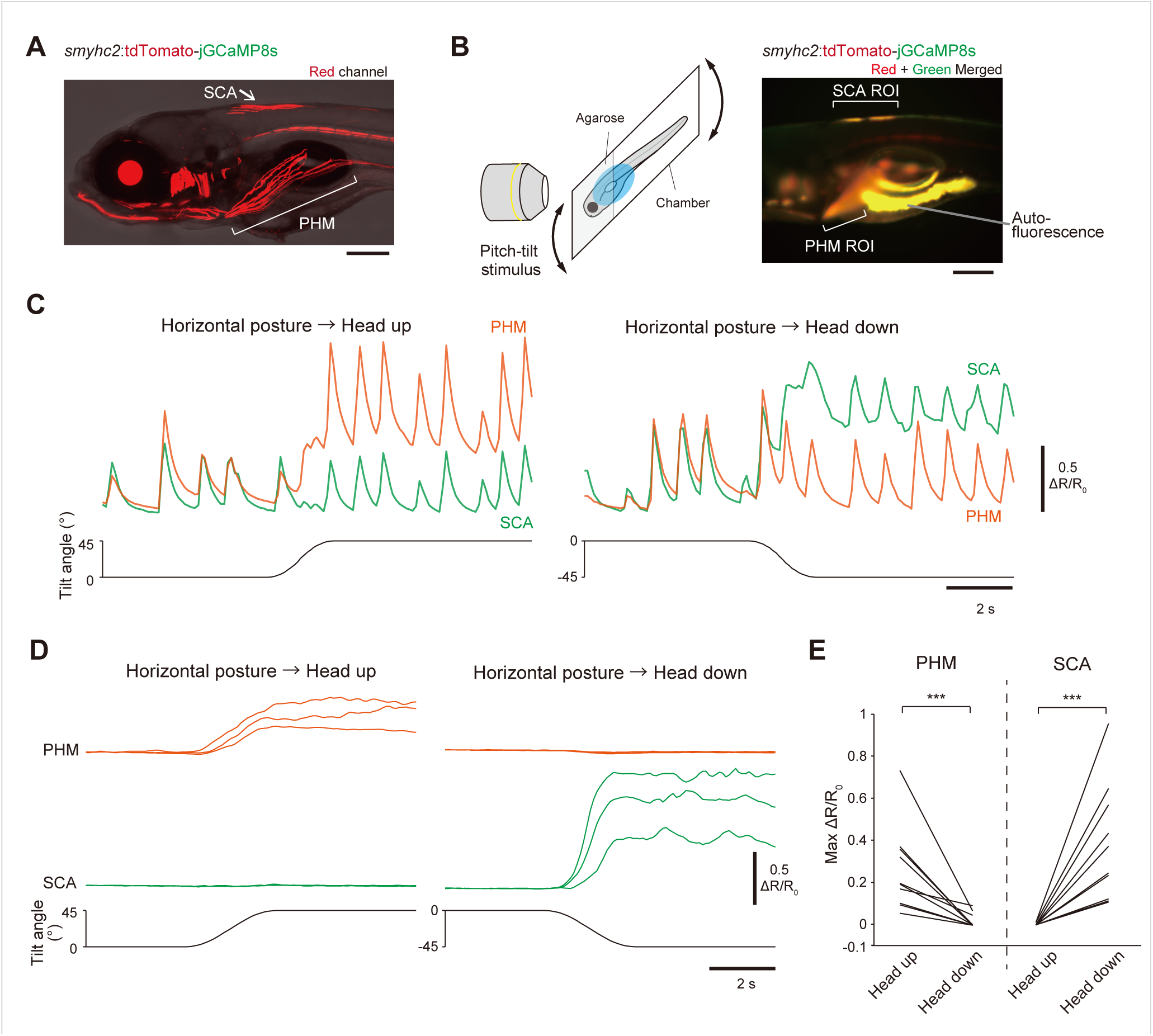
Ca^2+^ imaging of PHMs and SCAs during pitch tilts. (A) Confocal stack image of a Tg(*smyhc2*:tdTomato-jGCaMP8s) fish (maximum intensity projection). Red channel and transmitted light are merged. (B) Left: schematic of Ca^2+^ imaging with a tiltable-objective microscope. Right: fluorescence image of a Tg(*smyhc2*:tdTomato-jGCaMP8s) fish. Red and green channels are merged. Autofluorescence is visible in the intestine and yolk. Regions of interest (ROIs) covering the entire SCAs, and covering the rostral region of PHMs were simultaneously measured. (C) Ca^2+^ responses quantified as changes in green/red fluorescent intensity ratio (ΔR/R_0_) in PHMs (orange) and SCAs (green) in response to pitch tilts. Example traces showing sustained Ca^2+^ responses with fast Ca^2+^ transients. (D) Similar to (C), but under conditions with reduced fast Ca^2+^ transients. Three trials for each tilt stimulus condition from a single fish are shown. (E) Pairwise comparison of maximum ΔR/R_0_ values in PHMs and SCAs between head-up and head-down tilts under conditions with reduced fast Ca^2+^ transients (10 fish). For each stimulus, mean value of two to four trials is plotted. *p* = 4.3 x 10^-5^ (PHM), *p* = 1.1 x 10^-5^ (SCA) (Wilcoxon exact rank-sum test). Scale bars, 200 μm.

To examine whether *smyhc2*-positive PHMs and SCAs are involved in the ventral and dorsal flexions, we conducted Ca^2+^ imaging of these muscles during pitch tilts. The head of a Tg(*smyhc2*:tdTomato-jGCaMP8s) fish was embedded in agarose, and both PHMs and SCAs were simultaneously imaged from the lateral side using a custom-built tiltable-objective microscope^23,27^ (Figure 3B, left). SCAs were analyzed as entire regions, whereas PHMs were analyzed only in the rostral region because caudal regions overlapped with autofluorescence from the yolk and intestine (Figure 3B, right). Both PHM and SCA regions were analyzed by calculating ratiometric intensity change (ΔR/R_0_). During Ca^2+^ imaging, we frequently observed fast Ca^2+^ transients in PHMs and SCAs (Figure 3C), which coincided with jaw movements and Ca^2+^ transients in the lower jaw muscles (Figures S2A, S2B). We inferred that these fast Ca^2+^ transients were associated with mouth-opening movements, possibly related to respiration. Notably, in addition to these fast Ca^2+^ transients, a sustained increase in baseline activity was observed in PHMs during head-up tilt (orange trace in Figure 3C, left) and in SCAs during head-down tilt (green trace in Figure 3C, right). In contrast, neither SCAs during head-up tilt nor PHMs during head-down tilt showed such sustained increases (Figure 3C; green trace in the left panel; orange trace in the right panel). Thus, muscles on the flexed side were activated in response to pitch tilts.

To quantify muscle activities, we addressed the difficulty that tilt-induced Ca^2+^ responses were hard to quantify accurately, as fast Ca^2+^ transients often terminated abruptly or changed in frequency during tilts (Figure S2C). To mitigate this problem, we sought conditions that minimized fast Ca^2+^ transients and found suitable conditions in which the lower jaw of a fish was partially injured (see Methods). Figure 3D shows muscle activity under these conditions, in which tilt-induced responses were clearly observed and were quantifiable. ΔR/R_0_ values in muscles on the flexed side (PHMs during head-up tilts and SCAs during head-down tilts) showed a pronounced increase, whereas those on the stretched side (SCAs during head-up tilts and PHMs during head-down tilts) did not show significant changes (Figures 3D, S2D–S2F). Population data confirmed that ΔR/R_0_ values in muscles on the flexed side were significantly higher than those on the stretched side (Figure 3E, 10 fish).

We then investigated which regions of muscles were activated during pitch tilts. SCAs are composed of two segments (Figures 3B, right; S3A), and PHMs consist of three segments^27^. We measured ΔR/R_0_ values from both two SCA segments and found that both segments were active during head-down tilts (Figures S3C, S3D, and S3G). For the three PHM segments, Ca^2+^ imaging was performed from the ventral side to avoid autofluorescence overlapping the middle and caudal segments (Figure S3E). Results revealed that all PHM segments were active during head-up tilts (Figures S3F, S3H). Collectively, Ca^2+^ imaging revealed that *smyhc2*-expressing muscles on the flexed side (PHMs during head-up tilts and SCAs during head-down tilts) were activated during tilts, with all muscle segments on the flexed side responding to the tilts.

### PHMs and SCAs play important roles in the ventral and dorsal flexions of the body, respectively

We next asked whether *smyhc2*-expressing PHMs and SCAs are required for the ventral and dorsal flexions of the rostral body. To address this question, we bilaterally ablated either the PHMs or SCAs using genetic methods. Tg(*smyhc2*:loxP-RFP-loxP-DTA) fish were crossed with either Tg(*tbx2b*:Cre) for PHM ablation^27^, or Tg(*zic1*:Cre) for SCA ablation. Successful ablation was confirmed by immunostaining with the S58 antibody, which labels slow muscle fibers^31^. In Cre-negative fish, PHMs, SCAs, and axial muscles were positively stained (Figure 4A, top). In *tbx2b*:Cre-positive fish, labeling was specifically abolished in PHMs, while staining in SCAs and axial muscles remained (Figure 4A, middle). In contrast, in *zic1*:Cre-positive fish, staining in SCAs was absent, whereas PHMs and axial muscles appeared unaffected (Figure 4A, bottom).

**Figure 4.**
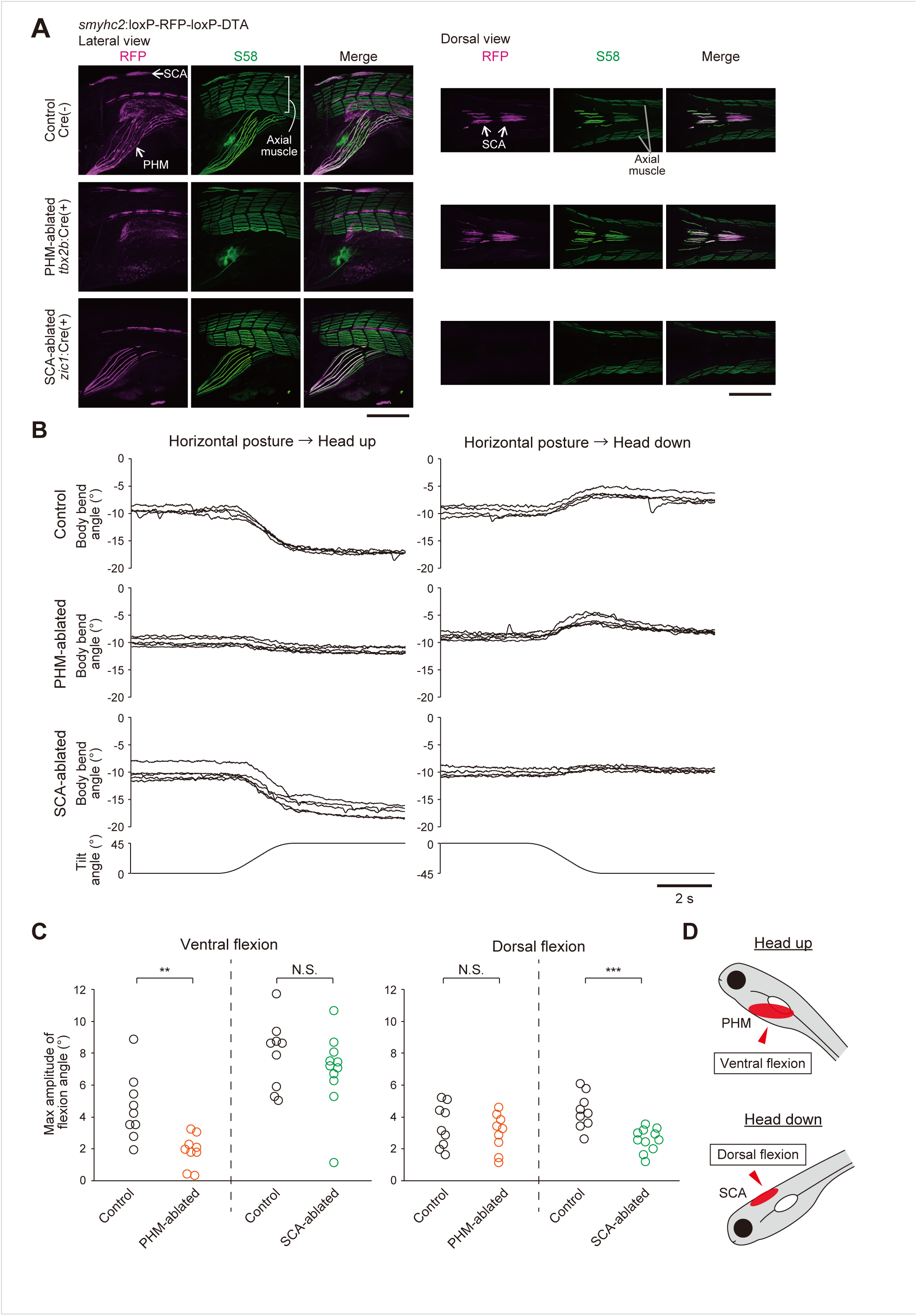
Body flexion in PHM- or SCA-ablated animals. (A) Immunohistochemistry images (confocal stacked images) of Tg(*smyhc2*:loxP-RFP-loxP-DTA) fish stained with S58 antibody. Images show fish without Cre (control, top row), with *tbx2b*:Cre (PHM-ablated, middle row), and with *zic1*:Cre (SCA-ablated, bottom row). Left three columns show lateral-view images, with rostral to the left and dorsal to the top. Right three columns show dorsal-view images, with rostral to the left. Scale bars, 200 μm. (B) Body bend angle traces of control (top), PHM-ablated (middle), and SCA-ablated fish (bottom) in response to pitch tilts under head-embedded conditions. For each group, trials from the same fish are shown. Four to five trials are shown for each condition (six conditions total). (C) Pairwise comparison of maximum amplitudes of ventral and dorsal flexion angles between control and muscle-ablated animals. Mean values of three to five trials are shown for each condition. Non-ablated (Cre-negative) siblings were used as controls. Orange: PHM-ablated fish (9 fish), green: SCA-ablated fish (11 fish), black: control groups (9 fish for each group). For ventral flexion: *p* = 0.001 (PHM ablation), *p* = 0.37 (SCA ablation); for dorsal flexion: *p* = 0.60 (PHM ablation), *p* = 0.0003 (SCA ablation) (Wilcoxon exact rank-sum test). (D) Summary of muscles involved in the body flexions. The ventral flexion during head-up tilts is produced by PHMs, whereas the dorsal flexion during head-down tilts is produced by SCAs.

We examined the effects of muscle ablation on the dorsal and ventral flexions in these animals under head-embedded conditions. Cre-negative control fish exhibited both types of flexions of the body (Figure 4B, top). PHM-ablated fish exhibited reduced ventral flexion (Figure 4B, middle left), while the dorsal flexion was retained (Figure 4B, middle right). SCA-ablated fish, on the other hand, exhibited reduced dorsal flexion (Figure 4B, bottom right), while the ventral flexion was retained (Figure 4B, bottom left). Data from multiple animals (9–11 fish) showed that ventral flexion angles were significantly smaller in PHM-ablated fish than in control siblings (orange circles in Figure 4C, left). Similarly, the dorsal flexion was significantly reduced in SCA-ablated fish (green circles in Figure 4C, right). Thus, the flexion toward the ablated side was impaired. In contrast, the flexion toward the intact side (ventral flexion in SCA-ablated fish and dorsal flexion in PHM-ablated fish) did not differ significantly from that of controls (Figure 4C; green circles in the left panel; orange circles in the right panel). These results demonstrate that *smyhc2*-expressing PHMs and SCAs play crucial roles in the ventral flexion and dorsal flexion, respectively (Figure 4D).

### The tangential nucleus plays a critical role in both dorsal and ventral flexions

We next investigated the neural circuits responsible for these two types of body flexions. It is known that tangential nucleus neurons (TAN neurons) in the vestibular nuclei receive utricular inputs that encode tilt orientation^23,32,33^, and that subset of them becomes active during head-up tilts^34^ (Figure 5A). Moreover, our previous study showed that vestibular information from TAN neurons is ultimately relayed to PHMs via the nucleus of the medial longitudinal fasciculus (nMLF), a class of reticulospinal neurons^27^. Therefore, we hypothesized that PHM-driven ventral flexion during head-up tilts is mediated by TAN–nMLF circuits. These circuits would be activated bilaterally, as pitch tilts elicit similar responses in both inner ears. The previous report also revealed that some TAN neurons are active during head-down tilts^34^ (Figure 5A). These data raise the possibility that TAN neurons in both left and right sides are also involved in the dorsal flexion during head-down tilts.

**Figure 5.**
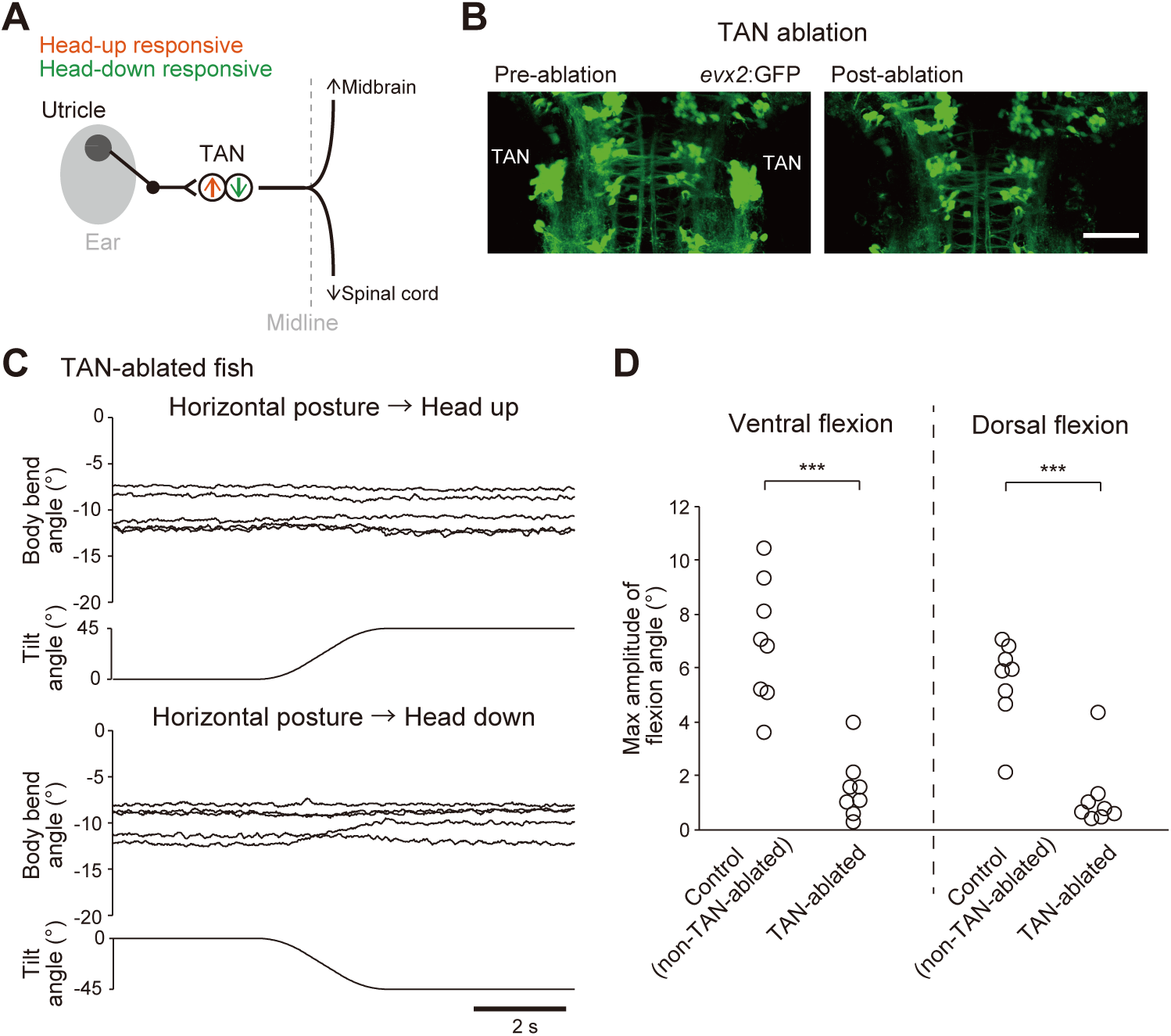
Body flexion in TAN-ablated fish. (A) Schematic of TAN neurons receiving utricular inputs. TAN neurons project their axons contralaterally and their branches extend rostrally into the midbrain and caudally into the spinal cord. Orange and green arrows indicate TAN neurons with response selectivity to head-up or head-down tilts. (B) Confocal stacked images of a Tg(*evx2*:GFP) fish before and after laser ablation of bilateral TAN neurons. Scale bar, 50 μm. (C) Body bend angle traces of a TAN-ablated fish in response to pitch tilts under head-embedded conditions. Five trials for each tilt stimulus condition from the same fish are shown. (D) Pairwise comparison of maximum amplitudes of ventral and dorsal flexion angles between control (non-TAN ablation; see Figure S4A and S4B) and TAN-ablated animals (8 fish each). Mean values of two to five trials are shown for each condition. *p* = 0.0003 (ventral flexion), *p* = 0.0003 (dorsal flexion) (Wilcoxon exact rank-sum test).

We tested whether TAN neurons are involved in the ventral and/or dorsal flexions by ablating TAN neurons bilaterally. The Tg(*evx2*:GFP) line labels excitatory TAN neurons, which project axons rostrally and caudally on the contralateral side^27,32,35^ (Figure 5A). We bilaterally ablated all GFP-positive TAN neurons (Figure 5B), and then measured flexion angles during pitch tilts under head-embedded conditions. As expected, the ventral flexion was impaired during head-up tilts (Figure 5C, top). The dorsal flexion was also abolished during head-down tilts (Figure 5C, bottom). As a control ablation experiment, we ablated *evx2*-positive non-TAN neurons (Figure S4A). Animals with non-TAN ablation showed no obvious impairment in either dorsal or ventral flexion (Figure S4B). Population data showed that both ventral and dorsal flexions in TAN-ablated fish were significantly smaller than those in non-TAN-ablated control fish (Figure 5D, 8 fish each). These results revealed that TAN neurons play an important role in both ventral and dorsal flexions of the body.

### Subsets of nMLF neurons are active during head-up and/or head-down tilts

We next focused on nMLF neurons, which receive inputs from TAN neurons^27,32^. We performed Ca^2+^ imaging of nMLF neurons during pitch tilts at single-cell resolution. These neurons were retrogradely labeled with the green fluorescent Ca^2+^ indicator Cal-520 dextran and the red fluorescent dye rhodamine dextran (Figure 6A). Some neurons (cells #1 and #2 in Figure 6B) increased ΔR/R_0_ values during head-down tilts, while others (cells #4–7) responded to head-up tilts. Some neurons, such as cell #3, responded to both tilts. Large soma-sized neurons (MeLr, MeLc, MeLm, and MeM) tended to exhibit little or relatively low ΔR/R_0_ values (cells #8–11). We analyzed a total of 266 neurons from 13 fish (192 neurons from 8 fish in Figure 6C and 74 neurons from 5 fish in Figure S5A). Neurons were considered active when ΔR/R_0_ exceeded 0.1, as in a previous study^23^. Of the 266 neurons, 97 neurons (36%) were active only during head-up tilts, while 26 neurons (10%) were active only during head-down tilts (Figures 6C, 6D, S5A). These neurons are collectively referred to as “direction-selective neurons”. Among the 110 neurons (41%) that were active during both tilts, approximately half of them (61 neurons) exhibited a ΔR/R_0_ value more than twice as high in one direction compared to the other (Figure 6D, gray dashed lines indicate a two-fold difference). These neurons are referred to as “direction-biased neurons”. Of the direction-selective or -biased neurons (97 + 26 + 61 = 184 neurons), the majority (143 neurons) were tuned to respond to head-up tilts, whereas the minority (41 neurons) were tuned to respond to head-down tilts. These two groups were spatially intermingled (Figure S5B, S5C). Together, these results suggest that nMLF neurons have partially overlapping but largely distinct populations for responding to head-up or head-down tilts, with a larger proportion responding to head-up tilts.

**Figure 6.**
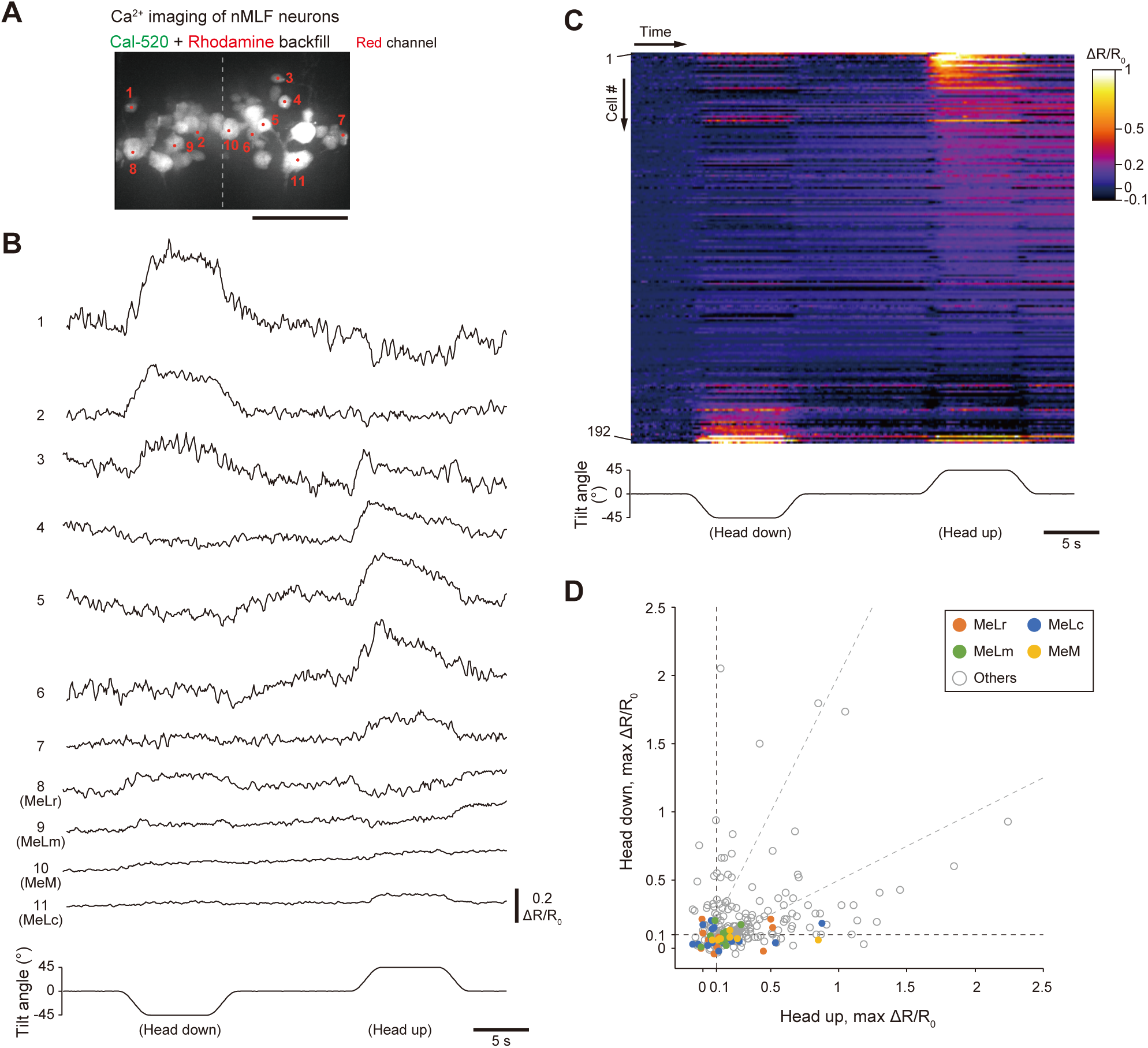
Ca^2+^ imaging of nMLF neurons. (A) Confocal stacked image of nMLF neurons retrogradely labeled with Ca^2+^ indicator Cal-520 and rhodamine. Only the red channel is shown. Dotted line indicates the midline. (B) Ca^2+^ responses of individual nMLF neurons in response to pitch tilts. Example response traces quantified as changes in green/red fluorescent intensity ratio (ΔR/R_0_). The numbers correspond to those shown in (A). (C) Color-coded ΔR/R_0_ time-series of 192 nMLF neurons from 8 fish in response to pitch tilts. Neurons are sorted based on the difference between head up/down tilt-induced ΔR/R_0_ amplitudes. See also Figure S5A. (D) Comparison of maximum ΔR/R_0_ values between head-up and head-down tilts for each neuron (total 226 neurons from 13 fish; 192 neurons from 8 fish in (C) and 74 neurons from 5 fish in Figure S5A). The black dashed lines indicate the artifact threshold of 0.1. The gray dashed lines indicate the boundaries for a two-fold difference.

### nMLF neurons play an important role in the ventral flexion during head-up tilts

To investigate whether nMLF neurons are necessary for both types of body flexions, we conducted behavioral assays with nMLF neuron function blocked. nMLF axons were bilaterally transected at rhombomere 1 using a laser in Tg(*nefma*:Gal4); Tg(UAS:GFP) fish (Figure 7A). The ascending axons of TAN neurons might also be transected, as they pass near the lesioned site. Figure 7B shows the body bend angle of the transected animals. The ventral flexion was abolished during head-up tilts (Figure 7B, top), whereas the dorsal flexion during head-down tilts was preserved (Figure 7B, bottom). As a control experiment, lateral longitudinal fascicles (LLF) fibers were bilaterally transected (Figure S6A). Animals with LLF transection showed no obvious impairment in either ventral or dorsal flexion (Figure S6B). In population data (obtained from 8–9 fish), ventral flexion angles in nMLF axon-transected fish were markedly reduced compared to those in the LLF-transected control group (Figure 7C, left). In contrast, the dorsal flexion agnles remained similar between the two groups (Figure 7C, right). These results suggest that nMLF neurons are necessary for the ventral flexion during head-up tilts, but not involved in the dorsal flexion during head-down tilts.

**Figure 7.**
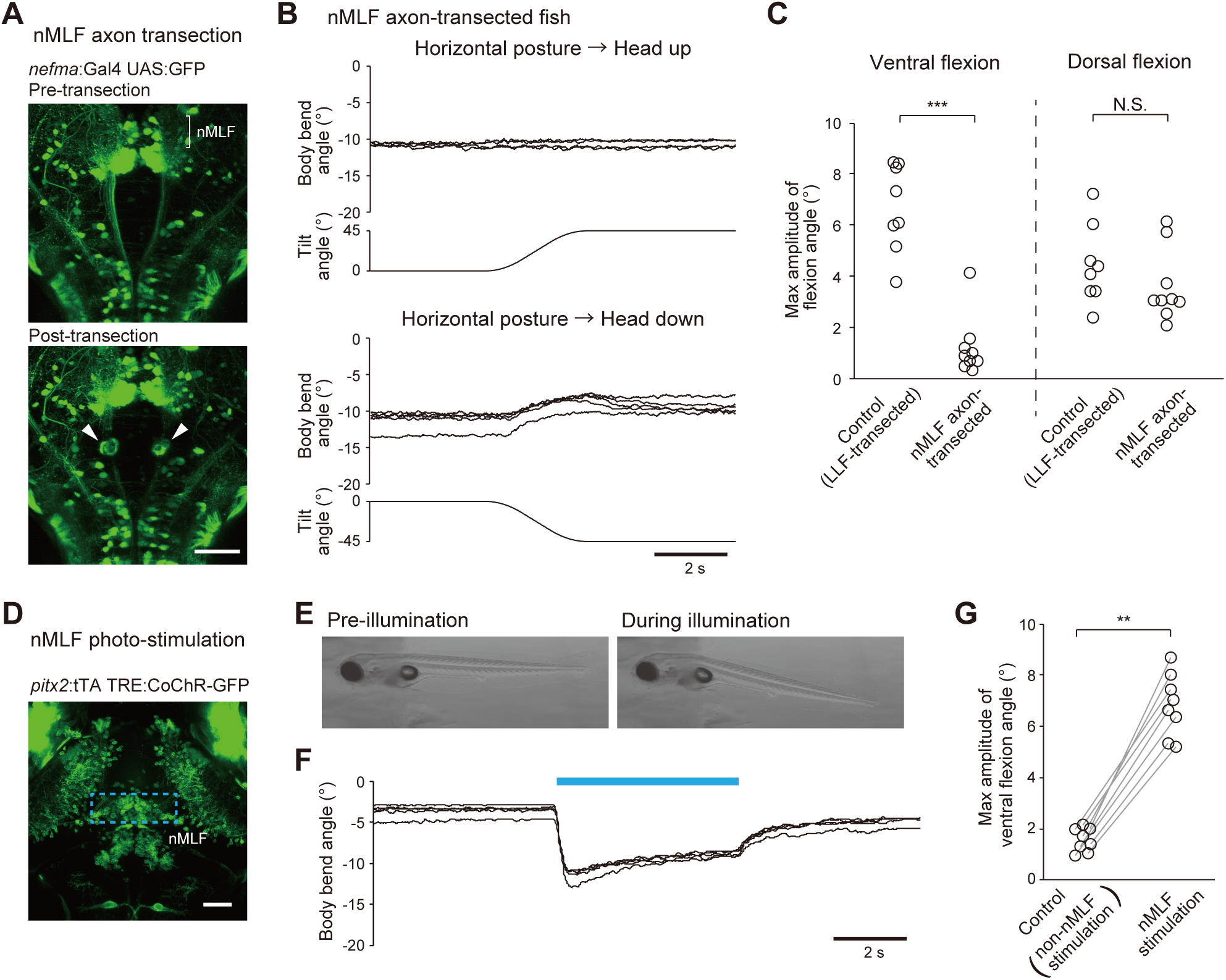
Axon transection and optogenetic activation of nMLF neurons. (A–C) Axon transection of nMLF neurons. (A) Confocal stacked images of a Tg(*nefma*:Gal4); Tg(UAS:GFP) fish before and after bilateral axon transection of nMLF neurons at rhombomere 1. (B) Body bend angle traces of a nMLF axon-transected fish in response to pitch tilts under head-embedded conditions. Four to five trials for each tilt stimulus condition from the same fish are shown. (C) Pairwise comparison of maximum amplitudes of ventral and dorsal flexion angles between control (LLF transection; 8 fish; see Figure S6A and S6B) and nMLF axon-transected animals (9 fish). Mean values of two to five trials are shown for each condition. *p* = 0.0001 (ventral flexion), *p* = 0.16 (dorsal flexion) (Wilcoxon exact rank-sum test). (D–G) Optogenetic activation of nMLF neurons. (D) Confocal stacked image of a Tg(*pitx2*:tTA); Tg(TRE:CoChR-GFP) fish. The blue dotted rectangle indicates the area illuminated with blue light. (E) Snapshots a fish before (left) and during (right) blue light illumination under head-embedded conditions. (F) Body bend angle traces upon blue light illumination (blue bar, 5 s). Five trials from a single fish are shown. (G) Pairwise comparison of maximum amplitude of ventral flexion angles between control (non-nMLF photo-stimulation; see Figure S6C and S6D) and nMLF photo-stimulation (8 fish). Both experiments were performed in the same fish. Mean values of four to five trials for each stimulation are shown. *p* = 0.007 (Wilcoxon signed-rank test). Scale bars, 50 μm.

In addition to the axon transection experiments, we performed optogenetic activation of bilateral nMLF neurons using Tg(*pitx2*:tTA); Tg(TRE:CoChR-GFP) (*pitx2* is expressed in approximately 80% of nMLF neurons^27^). Optogenetic activation of the bilateral nMLF neurons consistently elicited the ventral flexion (Figure 7D–7G; Movie S3). In a control experiment, activation of non-nMLF neurons elicited little flexion toward the ventral side (Figures 7G, S6C, S6D). Collectively, these ablation and optogenetic experiments indicate that nMLF neurons play a critical role in the ventral flexion during head-up tilts.

## Discussion

The vestibulo-collic reflex (VCR) is well characterized in tetrapod vertebrates^7–10^. Here, we first revealed that fish also exhibit a behavior resembling the VCR. Larval zebrafish precisely controlled the angle of the neck-like region depending on posture in the pitch axis: they flexed the region ventrally during head-up posture and dorsally during head-down posture. We further identified the neural circuits and muscles underlying these flexions. The ventral flexion during head-up posture was mediated by the TAN neurons through nMLF neurons and produced by PHMs. In contrast, the dorsal flexion during head-down tilts was also mediated by the TAN neurons, but did not require nMLF neurons, and was produced by SCAs (Figure 8; note that these circuits were recruited bilaterally).

**Figure 8.**
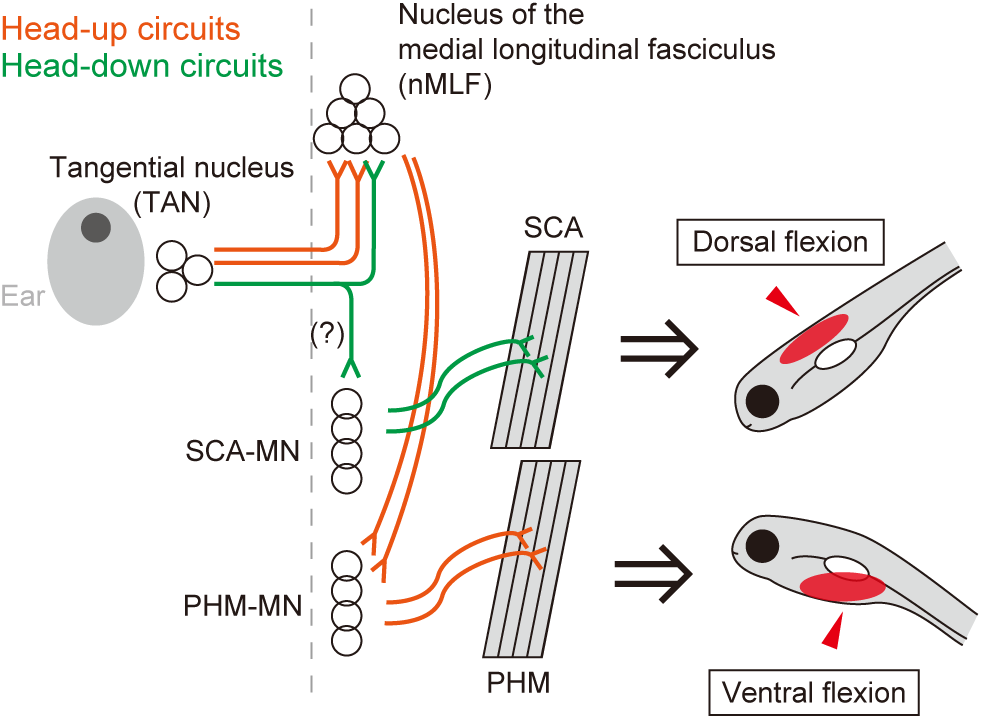
Circuit model for ventral and dorsal flexions of the body. During head-up tilts, the ventral flexion is mediated by TAN neurons and nMLF neurons, and produced by PHMs, which are located on the ventral side of the rostral body. During head-down tilts, the dorsal flexion is also mediated by TAN neurons, but likely bypassing nMLF neurons, and produced by the SCAs, which are located on the dorsal side of the rostral body. Note that the diagram shows only unilateral activation of the circuits; bilateral activation occurs *in vivo*. Rostral is to the top. The dashed line shows the midline.

### Contribution of the body flexion to head stabilization and movements

Our results showed that the body flexion helps bring the head closer to a horizontal orientation, partially contributing to the stabilization of the head orientation (Figure 2K). While mammals can achieve substantial head compensation through the VCR in response to vestibular stimuli^36,37^, larval zebrafish compensate only within a limited range of angles. This may be partly due to structural constraints, such as the absence of an anatomical neck. Despite this limited compensation, slight head stabilization together with vestibulo-ocular reflexes^32,38^ likely maintains visual focus and thereby enhances accurate spatial recognition, as observed in other animals^1,39^.

In addition to its role during pauses, the dorsal/ventral body flexion continues during swimming (Figure S1A), suggesting that it also functions during active movements. The body flexion alters the orientation of both head and tail (Figure 2), which likely influences the direction of locomotion. For example, adjusting head orientation toward the horizontal may help prevent excessive upward or downward deviation during escape behavior. During swimming, the dorsal/ventral body flexion may slightly modulate the direction of propulsive force generated by tail oscillations and contribute to active pitch control. Indeed, fish are known to adjust their pitch through swimming^29^, though the underlying biomechanisms are not fully understood. Further investigation on the body flexion may thus advance our understanding of fish postural control during swimming.

### Circuit mechanisms underlying dorsal and ventral flexions

Previous reports showed that distinct subsets of TAN neurons become active during head-up or head-down tilts, respectively^34^, and our results have shown that TAN neurons play an important role in both the ventral and dorsal flexions. However, the tilt information was transmitted through different pathways. The ventral flexion was mediated by nMLF neurons, presumably via ascending axons of TAN neurons, whereas the dorsal flexion did not require the nMLF neurons (Figure 8). What neuronal circuits mediate the dorsal flexion? One possibility is that descending branches of TAN neurons are involved in the dorsal flexion^27,32^ (Figure 5A). These branches extend into the rostral spinal cord^27^, where SCA motoneurons are located (Figure S3B). These results suggest that a subpopulation of TAN neurons may form monosynaptic connections to SCA motoneurons to produce the dorsal flexion. Notably, mammalian vestibular nucleus neurons directly connect to neck motoneurons^6^, and Xenopus TAN neurons directly connect to spinal motoneurons^40^. These studies support the possibility that monosynaptic connections underlie the dorsal flexion in fish.

If monosynaptic connections exist between TAN neurons and SCA motoneurons, the dorsal flexion circuit would consist of one fewer synapse than the ventral flexion circuit. The difference in circuit complexity between the dorsal and ventral flexions may reflect their functional requirements. We speculate that the ventral flexion may be modulated by a circuit with more synapses, including nMLF neurons, whereas the dorsal flexion is elicited in a more stereotyped manner by a circuit with fewer synapses. One example situation that requires modulation is prey capture. When prey is positioned slightly above, fish adopt a head-up posture and elevate their head^41^, which may require suppression of the vestibular-driven ventral flexion. This suggests that the ventral flexion may require context-dependent modulation through nMLF neurons, which integrate multiple inputs related to prey capture^41–43^. In contrast, fish constantly experience passive head-down torque^29^, and our data showed that they spent more time in head-down than in head-up postures (Figure 1C), making dorsal flexion the more frequently used response. Therefore, the dorsal flexion circuit may benefit from a simpler and more direct circuit that provides consistent responses.

nMLF neuronal population relay both head-up and head-down tilt information^25^ (Figure 6C). However, axon transection experiments revealed that nMLF neurons do not play a significant role in the dorsal flexion during head-down tilts (Figure 7B and 7C). We speculate that nMLF neurons also contribute to other aspects of pitch tilt responses. One possibility is that they modulate pectoral fin movements, as fins are finely coordinated with body movements during pitch tilts^44^. Further experiments will clarify the broader roles of nMLF neurons in tilt-induced behaviors.

### Pitch control vs. roll control

Larval zebrafish exhibit behavioral responses to roll and pitch tilts. Upon roll tilts, fish display a lateral body flexion toward the ear-up side^27^. This reflex repositions the animal’s center of gravity lateral to its center of buoyancy, generating a restoring torque that counteracts the roll. As a result, fish maintain a stable dorsal-up posture. In contrast, the dorsal or ventral flexion reflexes elicited by pitch tilts primarily function to stabilize the head rather than to maintain a horizontal body posture. This strategic difference likely reflects distinct functional demands. For roll control, larval zebrafish consistently maintain a dorsal-up posture^45^, underscoring its importance. In contrast, they display a wide range of pitch postures^46^ (Figure 1), indicating that precise horizontal alignment is less critical. These contrasting strategies—postural restoration for roll and head stabilization for pitch—highlight the axis-specific demands of orientation control in larval zebrafish.

At the neuronal circuit level, the TAN–nMLF–PHM pathway is engaged in both the ventral flexion during head-up tilts and the lateral body flexion during roll tilts. A key distinction lies in the laterality of the circuit activation: during roll tilts, unilateral activation of this circuit drives the lateral flexion to restore an upright posture, whereas during head-up pitch tilts, bilateral activation drives the ventral flexion. The difference between these two behaviors further lies in the different recruitment patterns of PHMs. During head-up tilts, all three PHM segments were activated in a similar manner (Figures S3F, S3H). In contrast, during roll tilts, all three PHM segments were active, but the increase in activity was smaller in the rostral segment than in the middle and caudal segments^27^. Thus, the rostral PHM segment appears to play a more prominent role in the ventral flexion compared to the lateral flexion. The rostral segment, located in the most ventral region of the PHMs, may be mechanically advantageous for producing the ventral flexion. In contrast, the middle and caudal segments, positioned just lateral to the swim bladder, may be better suited for the lateral flexion, as the lateral flexion near the swim bladder is thought to be the most energy-efficient behavior^27^. These results suggest that the ventral and lateral flexions are mediated by largely shared circuits, but at a finer level, the circuits function to increase efficiency for each movement. Together, these considerations highlight differences between roll and pitch control in fish.

An important question is how pitch and roll tilt information are integrated. TAN neurons receive utricular inputs that encode pitch and/or roll signals^23,33^. nMLF neurons, in turn, receive inputs from TAN neurons^27,32^, and are active in response to both pitch and roll tilts^22,25,27^ (Figure 6C). Thus, integration of pitch and roll information may occur at the TAN and/or nMLF levels. Further studies will elucidate the neuronal integration of the pitch and roll signals. In the natural environment, fish control their body in both pitch and roll axes simultaneously. The VCR-like behavior and self-righting movements presumably occur simultaneously to maintain head and body orientation.

### Evolutionary and comparative perspectives

Our results revealed that VCR-like behaviors are present in fish. Considering muscles and neural circuits, the body flexion of the neck-like region in zebrafish and the VCR in mammals exhibit conserved and divergent features across three aspects. We first focused on the muscles. Some mammalian VCR-related neck muscles (splenius capitis, suboccipital muscles) attach to the skull, while others (sternocleidomastoid, trapezius) attach to their pectoral girdle at the caudal ends^6,47^. In fish, SCAs of adult zebrafish attach to the skull^48^, consistent with their proposed function as “neck-like” muscles. However, PHMs attach to the pectoral girdle at their rostral ends^49^, representing a different anatomical arrangement from typical mammalian neck muscles. These features suggest that the body flexion in zebrafish recruits muscles that span a wider anatomical region compared with the discrete neck region in mammals, but both fish and mammals recruit muscles located immediately caudal to the head^50,51^ (Figure 3A).

Second, with respect to the vestibular nucleus, both reflexes share key mechanisms. Studies in mammals have shown that vestibular nucleus neurons connect to neck motoneurons both directly and indirectly^6,52^ (see below for the indirect pathway). Although it remains unclear which vestibular nucleus in mammals is analogous to zebrafish TAN, the projection pattern of TAN is similar to that of the mammalian vestibular nucleus: TAN neurons send neural signals to spinal motoneurons through nMLF neurons (indirect pathway) and through a potential direct pathway. However, there is a difference between mammalian and fish vestibular nuclei. In zebrafish, all TAN neurons project axons contralaterally^27,32^ (Figure 5A), whereas in mammals, vestibular neurons that directly connect to neck motoneurons project both contralaterally and ipsilaterally^53,54^.

Lastly, zebrafish nMLF neurons were recently proposed as analogous to the mammalian interstitial nucleus of Cajal (INC)^55^, which is involved in the VCR and eye movements^56,57^. Anatomically, both the INC and nMLF are located near the oculomotor nucleus in the midbrain^55,58^. Both nuclei receive inputs from the vestibular nuclei, visual system, and the cerebellum^6,21,22,52,56,59,60^. Both nuclei project to the spinal cord ipsilaterally and form monosynaptic connections to spinal motoneurons^56,61–63^. Our current study indicat that nMLF neurons are involved in head orientation, a feature similar to that of the INC. A difference between the two nuclei lies in their projection patterns. The INC projects to the spinal cord, reticular formation, extraocular motoneurons, and vestibular nuclei^62^. In contrast, nMLF neurons are known only to project to the spinal cord. However, these differences may reflect methodological factors in how the projections were identified: projections of the INC were traced by dye injection into the nucleus, whereas those of the nMLF were identified by retrograde labelling. Thus, there is a possibility that some neurons within the nMLF area project to additional targets, such as extraocular motoneurons. Discovery of such projections would strengthen the analogy between the nMLF and INC. Overall, although muscles and neural circuits differ to some extent, the zebrafish body flexion exhibits several features in common with the mammalian VCR.

Our findings suggest that the body flexion observed in zebrafish may represent a primitive form of the VCR, and that the VCR and its underlying circuits may be evolutionarily conserved across vertebrates. Our results further suggest that VCR circuits may have evolved before the emergence of an anatomical neck. In this sense, the body flexion in the neck-like region in zebrafish will serve as a useful model for investigating the fundamental mechanisms of vertebrate head stabilization and provide insights into the evolutionary origin of head-stabilizing behaviors.

## Methods

### Animals

All procedures were conducted in accordance with the guidelines approved by the Animal Care and Use Committees at the National Institute of Natural Sciences. Zebrafish adults, embryos, and larvae were maintained at 28.5 °C under a 14/10-hour light/dark cycle, except for embryos and larvae expressing channelrhodopsin, which were kept in darkness. All experiments were performed using 5–6 days post-fertilization (dpf) larvae. Sex was indeterminate at the larval stage.

The following previously published transgenic lines were used: Tg(*smyhc2*-hs:loxP-RFP-loxP-DTA)^27^, Tg(*tbx2b*-hs:Cre)^27^, Tg(*evx2*-hs:GFP)^64^, Tg(*nefma*-hs:Gal4);Tg(UAS:GFP)^65^, Tg(TRE:CoChR-GFP)^66^, Tg(*pitx2*-hs:Dendra2)^27^, and Tg(*actc1b*:GFP)^67^. The following transgenic lines were generated using the CRISPR/Cas9-mediated knock-in method^68^: Tg(*smyhc2*-hs:tdTomato-jGCaMP8s), Tg(*zic1*-hs:Cre), and Tg(*pitx2*-hs:tTA). Additionally, Tg(*smyhc2*:loxP-RFP-loxP-DTA) was generated, which lack heat shock promoter (hsp70). The sgRNA sequences for gene targeting were as follows: gacttggatttcatctggcg (for *smyhc2*), gcagcacgtgaccccgtgct (for *zic1*), and gagctttgactgtcagcgcg (for *pitx2*)^27^. Zebrafish lines generated in this study have been deposited to the National BioResource Project in Japan. Note that “-hs” is omitted in the Results and Figures/Figure legends.

### Behavioral recording of freely swimming fish

A 6-dpf larva was transferred to a thin vertical chamber (5.5 mm width × 2.5 mm height × 1 mm depth) constructed from two microscope glass slides and filled with fish-rearing water (Figure 1A). A digital camera (Teledyne FLIR, GS3-U3-23S6M), a 10-mm close-up extension tube (Tamron, MR-set), a lens (Tamron, M118FM50), and software (Teledyne FLIR, FlyCapture2) were used to capture the behavior at 100 frames per second (fps). Infrared light (CCS, LDL-60X60IR2-850) was illuminated from behind the chamber. Recording was manually initiated and terminated when the fish entered or exited the recording field of view (20 mm width × 13 mm height) in the center of the chamber.

### Behavioral experiments with agarose-embedded fish

To perform head-restrained behavioral experiments, a 6-dpf larva was anesthetized in 0.02% tricaine and was embedded lateral side down in 2% low-melting point agarose (Nacalai Tesque, #01161-12) in a small acrylic chamber (12 mm length × 15 mm width × 4 mm height). Agarose located posterior to the heart (ventral side) or the otic vesicles (dorsal side) was removed (Figure 2A; 2B, left). The chamber was filled with fish-rearing water and covered with a cover slip. The chamber was then placed vertically on a vertically mounted motorized rotation stage (Thorlabs, DDR100/M), so that the fish was imaged from the lateral side. Fish behavior was captured as described for the freely swimming fish, except that a longer close-up extension tube (20 mm, Tamron, MR-set) was used. Fast, rhythmic trunk movements were often observed immediately after the operation (Figure S1D). To suppress these fast movements, the fish was kept in the dorsal-up position for 10 to 30 minutes to acclimate to the embedded condition. Pitch-tilt stimuli of ±45° (maximum velocity, 25°/s; maximum acceleration and deceleration, 25°/s^2^) were then applied in the head-up or head-down directions.

The body-restrained fish preparation was similar to the head-restrained preparation described above, except for the following modifications. The agarose located anterior to the caudal end of the swim bladder was removed, and a fine tungsten pin was inserted into the notochord of the tail to anchor the fish in the agarose (Figure 2F; 2G, left).

### Ca^2+^ imaging

Ca^2+^ imaging was performed as previously described^23,27^. Briefly, ratiometric Ca^2+^ imaging was carried out using a tiltable objective microscope. An objective lens and a fish were tilted together using a vertically mounted motorized rotation stage (Thorlabs, DDR100/M). To image fish from the dorsal or ventral side, a 45° mirror was placed between the rotation stage and the objective lens^27,64^, while lateral-view images were obtained using the objective lens mounted directly on the rotation stage. The tilt stimuli were the same as those used in the head-restrained behavioral experiments. Blue excitation light was delivered to the sample, and green (jGCaMP8s or Cal-520 dexstran, Ca^2+^-dependent) and red (tdTomato or tetramethylrhodamine dextran) signals were simultaneously recorded using an image-splitting optics (Hamamatsu Photonics, W-View Gemini) and a single digital camera at 10 fps (Hamamatsu Photonics, ORCA-Flash4.0 V3).

For muscle imaging, we used 6-dpf Tg(*smyhc2*-hs:tdTomato-jGCaMP8s) fish with a *casper* background (*mitfa* ^-/-^; *roy*^-/-^)^69^. The fast Ca^2+^ transients of SCAs and PHMs were often observed in synchrony with jaw movements under head-restrained conditions (Figure S2B). To obtain tilt-induced Ca^2+^ responses with reduced fast Ca^2+^ transients, a fish was anesthetized in 0.02% tricaine and the lower jaw was partially injured. The lower jaw was stabbed approximately ten times with a tungsten pin. The fish was then head-embedded in a chamber, and covered with a cover slip. Muscle imaging was conducted using the widefield version of a tiltable objective microscope^27^ with a 10×/NA 0.3 objective lens (Olympus, UPLANFLN) and blue-light illumination (X-Cite, exacte, 3.0 mW/mm^2^ at sample). For simultaneous imaging of SCAs and PHMs, as well as for the imaging of segmental SCAs, muscles were imaged from the lateral side (Figures 3C–3E, S3C, S3D, S3G), whereas imaging of segmental PHMs was performed from the ventral side (Figure S3E, S3F, S3H).

To image nMLF neurons, the tip of a tungsten pin was immersed in a mixture of 25% (w/v) Cal-520 dextran and 25% (w/v) tetramethylrhodamine dextran and inserted into the spinal cord at the level of the second to fourth muscle segments in a 5-dpf nacre (*mitfa^-/-^*) fish^70^. The larva was allowed to recover until 6 dpf, then embedded in 2% low-melting point agarose (Nacalai Tesque, #01161-12), filled with fish-rearing water, and covered with a fluorinated ethylene propylene (FEP) sheet. Imaging was conducted using the confocal version of a tiltable objective microscope equipped with a spinning-disk confocal scanner (Yokogawa, CSU-X1), and a 40×/NA 0.8 water-immersion objective lens (Olympus, LUMPLANFLN)^23^. A 488-nm laser (COHERENT, Sapphire 488-50 CDRH) was used as the light source, with a laser power of 0.3–1.5 mW/mm^2^ at the sample. The first imaging series was performed in the most dorsal (or ventral) single optical plane, and subsequent series were performed sequentially toward more ventral (or dorsal) planes. After Ca^2+^ imaging, confocal images were acquired to determine soma positions using an inverted microscope (Leica Microsystems, TCS SP8 MP) equipped with a 40×/NA 1.10 objective lens (Leica Microsystems; #11506357).

### Evaluation of artifacts in a widefield version of tiltable objective microscope

Tiltable objective microscopes induce artificial changes in the fluorescence intensity arising from the tilt of a sample and muscle displacement. The artifact level associated with imaging fish from the dorsal or ventral side (using a tiltable microscope with a 45° mirror) was measured previously^27^ (confined within ±0.1). To assess artifact level when imaging fish from the lateral side (using an objective lens directly attached to the rotation stage), we used a 6-dpf Tg(*pitx2*-hs:Dendra2) fish that express Dendra2 at SCAs and PHMs. The fish was illuminated with violet light (400–420 nm) for 10 minutes to partially photoconvert green Dendra2 to red Dendra2 (Figure S3D). Imaging analysis of the SCAs and PHMs during ±45° tilt stimulatioin revealed that the amplitude of ΔR/R_0_ was confined within ±0.06 (Figure S2E).

### Muscle ablation

To genetically ablate muscles, we expressed the diphtheria toxin A fragment in a muscle-specific manner. Tg(*tbx2b*-hs:Cre); Tg(*smyhc2*-hs:loxP-RFP-loxP-DTA) fish was used for PHM ablation^64^: *tbx2b* is expressed in migratory muscle precursors, including those that give rise to PHMs^71^. Tg(*zic1*-hs:Cre); Tg(*smyhc2*:loxP-RFP-loxP-DTA) fish was used for SCA ablation: *zic1* is expressed in the dorsal part of somites^72^. Note that the reporter/toxin line used for PHM ablation, Tg(*smyhc2*-hs:loxP-RFP-loxP-DTA), contains heat shock promoter (hsp70), whereas the line used for SCA ablation, Tg(*smyhc2*:loxP-RFP-loxP-DTA), does not contain hsp70. This is because leaky reporter expression was subtly observed in axial muscles in Tg(*smyhc2*-hs:loxP-RFP-loxP-DTA). When crossed with Tg(*zic1*-hs:Cre), the axial epaxial muscles was partially ablated in addition to SCAs. To prevent this unwanted ablation, we created Tg(*smyhc2*:loxP-RFP-loxP-DTA), in which reporter/toxin expression was restricted to SCAs (Figure 4A).

### Immunohistochemistry

Immunostaining was performed using 6-dpf fish according to Doganli et al.^73^. The S58 monoclonal antibody supernatant (primary antibody; DSHB, RRID: AB_528377), which labels slow muscle fibers^31^, was used at a 1:10 dilution. Alexa Fluor 488-conjugated goat anti-mouse IgG secondary antibody (Thermo Fisher Scientific, RRID: AB_2534088) was used at a 1:500 dilution.

### Laser ablation

Laser ablation of TAN neurons was performed in 5-dpf Tg(*evx2*-hs:GFP) larvae. A larva was anesthetized in 0.02% tricaine and embedded in 1.5% low-melting point agarose in the dorsal-down orientation. The fish was then placed on an inverted microscope (Leica Microsystems, TCS SP8 MP). Ablation was performed using a two-photon laser (900 nm; InSight DeepSee 680–1300 nm; 52 mW/mm^2^ at the sample) and a 40×/NA 1.10 objective lens (Leica Microsystems; #11506352). The field of view was zoomed to approximately the size of a single cell, and 2–3 scans were applied to ablate the neuron. Bilateral GFP-labeled TAN neurons were subjected to laser ablation. Afterward, the larva was allowed to recover and was assayed at 6 dpf. For control ablation experiments, GFP-positive neurons located on the lateral side of rhombomere 4 were subjected to laser ablation.

Axon transection of nMLF neurons was carried out using Tg(*nefma*-hs:GFP); Tg(UAS:GFP) larvae in the same manner as the ablation of TAN neurons. Bilateral nMLF axons were severed at rhombomere 1. For control experiments, bilateral LLFs were transected at rhombomere 1.

### Optogenetic activation experiments

To optogenetically activate nMLF neurons, a Tg(*pitx2*-hs:tTA);Tg(TRE:CoChR-GFP) fish at 6-dpf was head-embedded dorsal side up in 2% low-melting point agarose in an acrylic chamber equipped with a glass cover slip on the lateral side. Agarose located caudal to the swim bladder was removed. The chamber was placed under an upright microscope (Olympus, BX51WI) equipped with a 10×/NA 0.3 water immersion objective lens (Olympus, UMPLANFLN). Behavior was imaged from the cover slip using a camera, lens, and extension tube, as described for the agarose-embedded experiments. nMLF neurons in a rectangular region (150 μm × 50 μm) were selectively illuminated with blue LED light (Prizmatix, UHP-LED-470, 3.2 mW/mm^2^ at sample) for 5 seconds using a digital mirror device (Mightex, Polygon400). For the control illumination experiment, *pitx2*-positive neurons located rostral to the eyes were illuminated in the same animal.

### Behavioral analysis

Behavioral analyses were performed using custom-written Python scripts based on OpenCV. Speed, pitch angle, and rostral body bend angle were calculated for each frame. Speed was calculated from the displacement of the center of the swim bladder. A pause was defined as a period when the speed was less than 1 mm/s, and swimming was defined as a period when the speed exceeded 5 mm/s (the same as a previous study^29^). Pauses longer than 0.2 seconds and swimming bouts longer than 0.05 seconds were analyzed. The pitch angle was defined as the angle between the horizontal line and a line connecting the center of the eye and the center of the swim bladder. The body bend angle was defined as the angle between lines fitted to the head and trunk contours as follows. Images were rotated, so that the fish faced leftward and pitch angle in the rotated image becomes zero. The rotated images were binarized and the dorsal body head and trunk contours were detected. The head contour line was fitted by connecting the dorsal contours at the level of the swim bladder and that at the level of 0.4 mm rostral to the swim bladder. The trunk contour line was fitted by connecting the dorsal contours at the level of 0.4 and 0.8 mm caudal to the swim bladder.

In behavioral analysis of freely moving fish, the pitch and body bend angles are prone to misestimation when the fish slightly yawed relative to the camera (i.e., not perfectly perpendicular to the viewing axis). To assess yawing, we also measured apparent eye size, which increases when the contralateral eye becomes visible as the fish yaws, and the total length of the dorsal contour from the head tip to the tail end, which decreases during yawing. If the detected eye size exceeded 1.5 times its minimal value, or if the total length of the dorsal contour fell below 0.8 times its maximum value, the corresponding frame was excluded from analysis. Additionally, periods in which the fish exhibited extreme pitch postures (pitch angle > 60° or < -60°) were excluded from analysis.

In the analysis of head- or body-embedded behavioral experiments, trials involving swimming or fast rhythmic trunk movements (Figure S1D) were excluded. A three-frame moving average was applied to smoothen the body bend angle time course. The body bend angles during the horizontal posture under restrained conditions were more negative (approximately –10°; Figures 2D, 2I, S1C) than those observed under freely moving conditions (approximately –5°; Figure 1C). This is likely due to the agarose embedding procedure. The flexion angle was defined as the change in the body bend angle during tilted position relative to that during the pre-stimulus horizontal position. The maximum amplitude of the flexion angle was calculated for each trial. In Figure 2E and S1C, wild-type animals and Cre-negative control groups in the muscle-ablation experiments were pooled.

### Analysis of Ca^2+^ imaging

Imaging analyses were performed using custom written Python scripts based on OpenCV and ImageJ. Images recorded by the tiltable-objective microscopes rotated during tilt stimulus. For image registration purposes, green and red channel images were counter-rotated. The images were then registered in the xy direction to correct minor image translation. Region of interests were manually drawn in ImageJ. The ΔR/R_0_ values of the region of interests in each frame were calculated, where R_0_ was the average ratio of the ten frames preceding tilt stimulus. A three-frame moving average was applied and the maximum ΔR/R_0_ value was then calculated.

The positions of the nMLF neurons recorded from different fish were normalized relative to the midline (for medio-lateral axis) and the position of MeLm neurons (for rostro-caudal and dorso-ventral axes). When both left and right MeLm neurons were labeled, the normalized rostro-caudal position was referenced to more caudal MeLm neuron, and the normalized dorso-ventral axis position was referenced to the ipsilateral MeLm neuron.

### Statistical analysis

For most of the experiments, multiple trials were conducted in a single fish. In Ca^2+^ imaging of nMLF neurons, neurons in the same optical plane could not be imaged repeatedly due to the desensitization of Cal-520. In experiments with multiple trials in a single fish, the mean value was calculated, and used as a representative result for that fish. Statistical analysis was performed using R. Statistical significance was assessed with the two-tailed Wilcoxon exact rank-sum test or two-tailed Wilcoxon signed-rank test. Statistical results were indicated as follows: *, *p*-value <0.05; **, *p*-value <0.01; ***, *p*-value <0.001; or N.S. (not significant), *p*-value ≥0.05.

## Data, code, and material availability

Further information and requests for resources, reagents, data, and codes are available from Shin-ichi Higashijima. Zebrafish lines generated in this study have been deposited to the National BioResource Project in Japan.

## Acknowledgements

We thank Higashijima lab members for their help with generating transgenic fish, fish care, and discussion; and the Optics and Imaging Facility of the National Institute for Basic Biology for the use of their microscopes. This work was supported in part by grants from the Ministry of Education, Culture, Sports, Science and Technology of Japan and from National Bioresource Project in Japan (KAKENHI Grant Numbers JP24KJ0242, JP24K18163 to T.S., JP20K06866, JP23K05983 to M.T., JP23K23929, JP25K02315 to S.H.) and MEXT National BioResource Project (NBRP) (to S.H.).

## Author Contributions

Conceptualization, T.S., M.T., and S.H.; Methodology, T.S., T.R.T., H.B., M.T., and S.H.; Investigation, T.S., T.R.T.; Software, T.S.; Formal Analysis, T.S.; Data Curation, T.S.; Writing – Original Draft, T.S.; Writing –Review & Editing, T.S., T.R.T., H.B., M.T., and S.H.; Visualization, T.S.; Supervision, M.T., and S.H.; Project Administration, S.H.; Funding Acquisition, T.S., M.T., and S.H.

## Declaration of generative AI and AI-assisted technologies in the writing process

During the preparation of this work, the authors used ChatGPT to improve the readability. After using this tool, the authors reviewed and edited the content as needed and take full responsibility for the content of the publication.

## Figure legends

**Figure S1.**
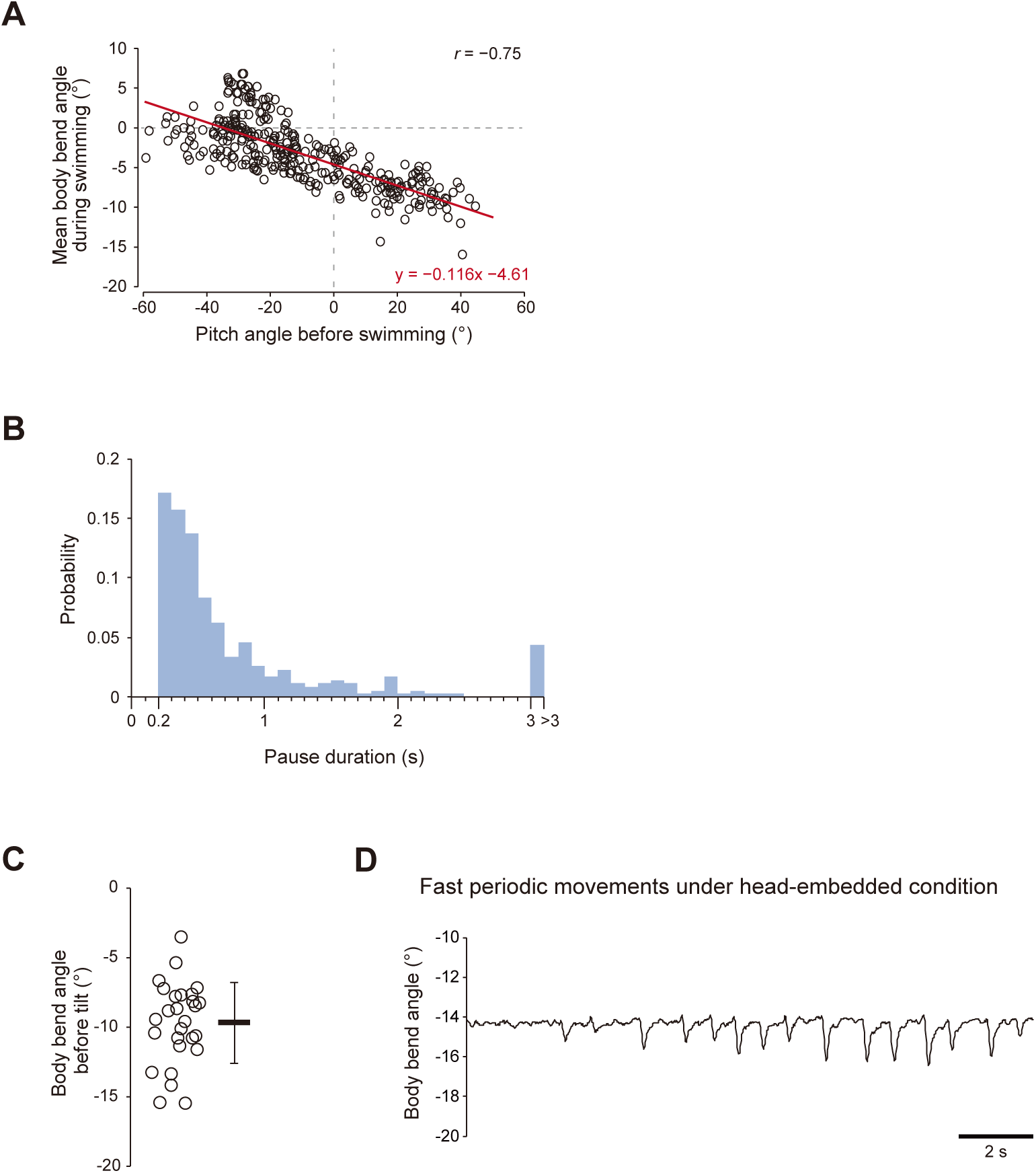
Behavior of freely moving fish and head-restrained fish. (A) Body bend angles during swimming plotted against the pitch angles immediately before swimming onset. Swimming bouts longer than 0.05 seconds were analyzed (334 swimming bouts from 12 fish). A regression line is shown in red. *r* represents Pearson correlation coefficient. (B) Distribution of pause duration. Pauses longer than 0.2 seconds were analyzed (340 pauses from 12 fish), and those exceeding 3 seconds were grouped into a single bin. (C) Body bend angle before tilt stimuli under head-embedded conditions (27 fish). For each fish, the mean value of four to ten trials is plotted. Thick line indicates the mean, and thin lines represent standard deviation. (D) Body bend angle trace showing fast periodic trunk movements under head-embedded conditions.

**Figure S2.**
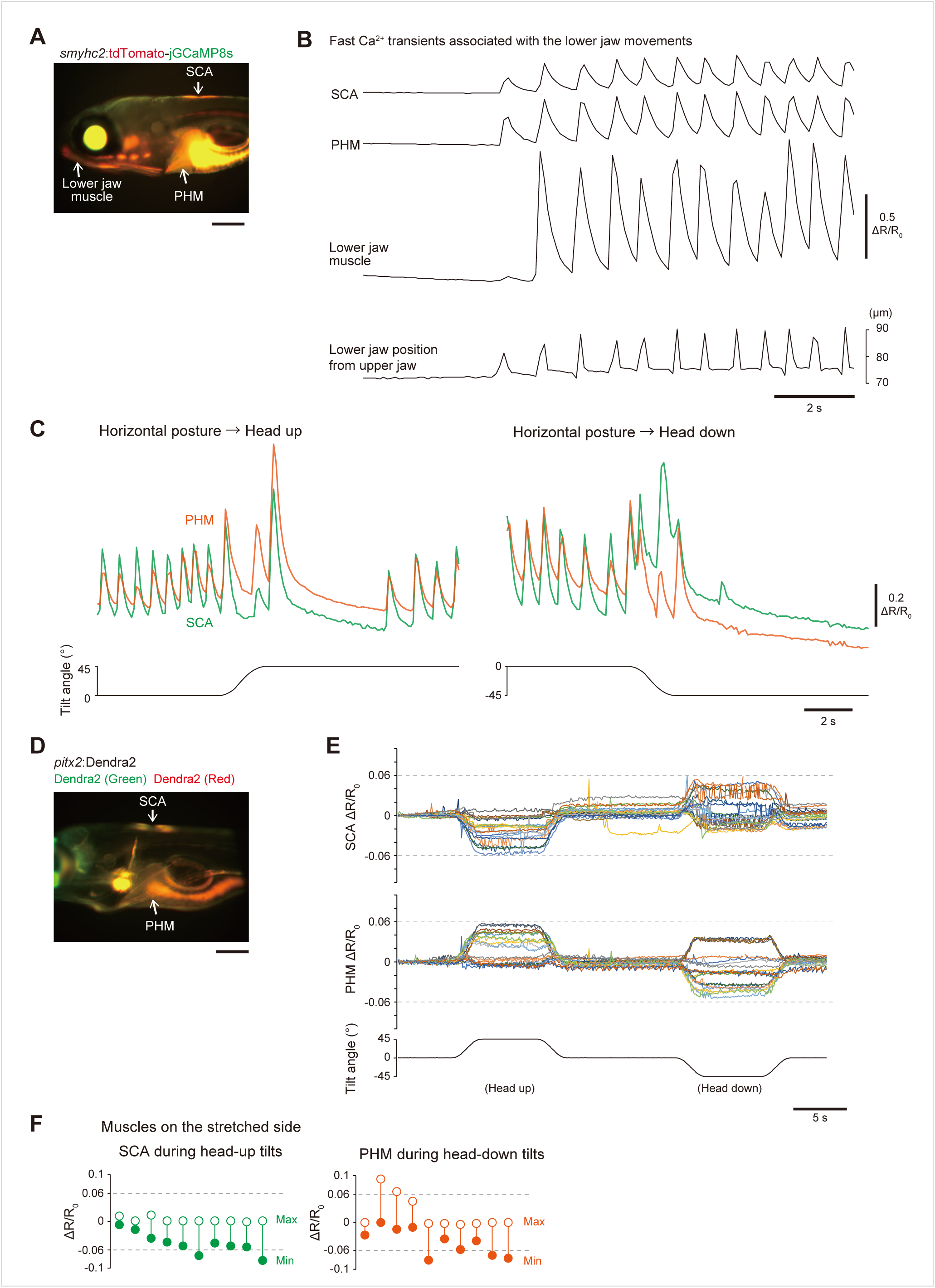
Ca^2+^ imaging of SCAs and PHMs, and characterization of artifact level. (A–C) Fast Ca^2+^ transients of SCAs and PHMs. (A) Lateral-view image of a Tg(*smyhc2*:tdTomato-jGCaMP8s) fish. (B) Example Ca^2+^ responses of SCAs, PHMs, and lower jaw muscles, and the position of the lower jaw, under head-embedded conditions. Ca^2+^ response were quantified as changes in green/red fluorescent intensity ratio (ΔR/R_0_). (C) Example ΔR/R_0_ traces of SCAs and PHMs in response to pitch tilts, showing termination or resumption of fast Ca^2+^ transients. (D, E) Characterization of an artificial ΔR/R_0_ level of the tiltable-objective microscope. (D) Lateral-view image of a Tg(*pitx2*:Dendra2) fish after a partial photoconversion of Dendra2. *pitx2* is expressed at SCAs and PHMs. (E) ΔR/R_0_ traces of SCAs and PHMs. All 21 trials from 6 fish are shown. Each color represents an individual trial. Tilt-driven artifacts remained within ±0.06, shown as dashed lines. (F) Maximum and minimum ΔR/R_0_ values in muscles on the stretched side under conditions with reduced fast Ca^2+^ transients (10 fish). Left: SCAs during head-up tilts; right: PHMs during head-down tilts. Open circles represent maximum values, and closed circles represent minimum values. Each pair from the same fish is connected by a vertical line. Gray dashed lines indicate artifact threshold of ±0.06. Scale bars, 200 μm.

**Figure S3.**
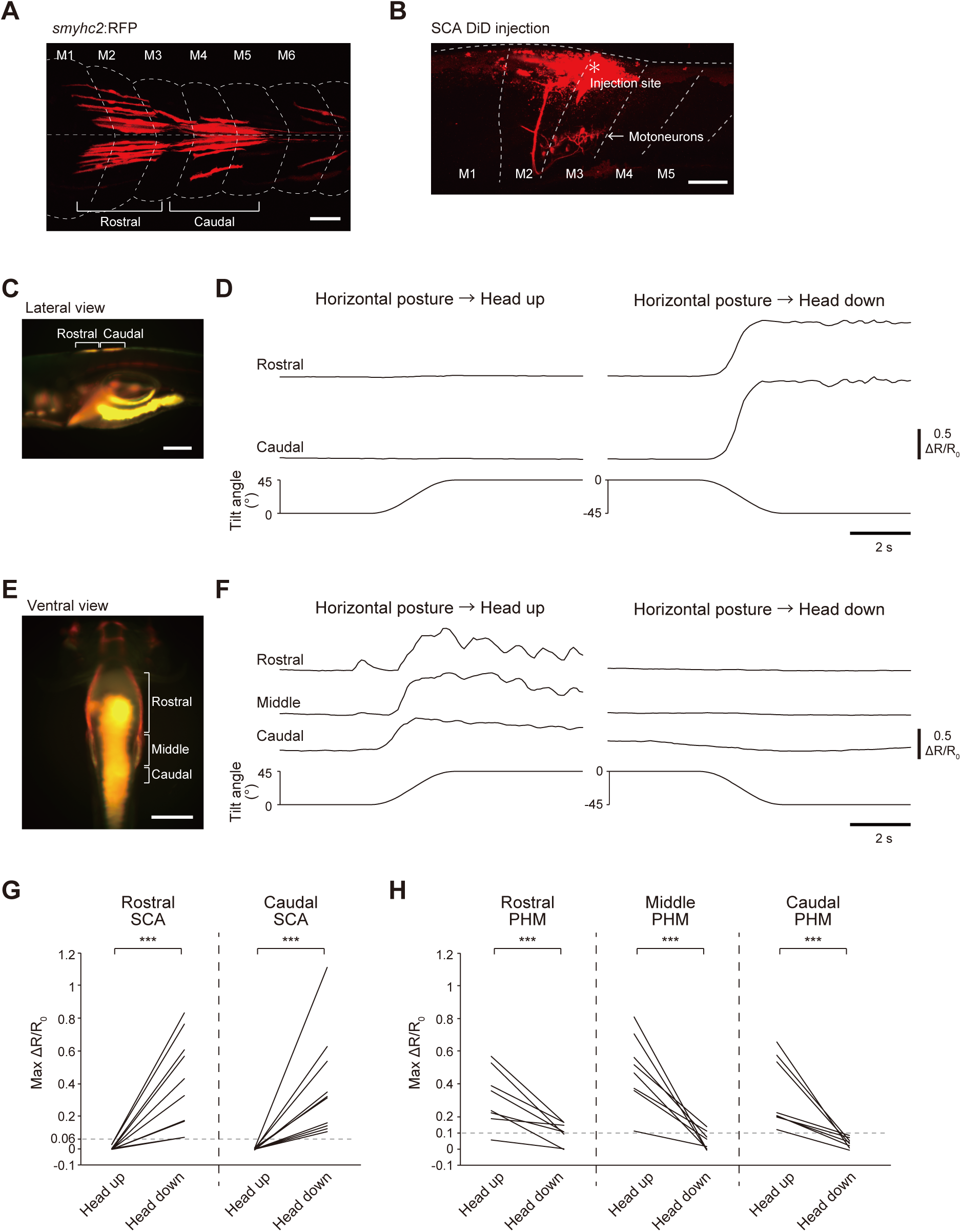
**Anatomical characterization of SCAs, SCA motoneurons, and segmental Ca^2+^ imaging of muscles** (B) Lateral-view image of retrograde labelling of SCA motoneurons. Dashed lines indicate the boundaries of the body or muscle segments. Rostral is to the left. Dorsal is to the top. (C, D) Ca^2+^ imaging of SCAs in each muscle segment. (C) Lateral-view image of a Tg(*smyhc2*:tdTomato-jGCaMP8s) fish. Rostral is to the left. Dorsal is to the top. (D) Example Ca^2+^ responses of rostral and caudal segments of SCAs in response to pitch tilts under conditions with reduced Ca^2+^ transients. Ca^2+^ response were quantified as changes in green/red fluorescent intensity ratio (ΔR/R_0_). (E, F) Ca^2+^ imaging of PHMs in each muscle segment. (E) Ventral-view image of a Tg(*smyhc2*:tdTomato-jGCaMP8s) fish. Rostral is to the top. (F) Example ΔR/R_0_ traces of rostral, middle, and caudal segments of PHMs in response to pitch tilts under conditions with reduced Ca^2+^ transients. (G) Pairwise comparison of maximum ΔR/R_0_ values in each muscle segment of SCAs between head-up and head-down tilts under conditions with reduced fast Ca^2+^ transients (10 fish). Mean values of two to four trials are shown for each tilt stimulus condition. *p* = 1.1 x 10^-5^ (rostral), *p* = 1.1 x 10^-5^ (caudal) (Wilcoxon exact rank-sum test). Gray dashed lines indicate the artifact threshold of 0.06. (H) Same as (G), but for each muscle segment of PHMs (8 fish). Mean values of four to five trials are shown for each tilt stimulus condition. *p* = 0.004 (rostral), *p* = 0.00006 (middle), *p* = 0.0001 (caudal) (Wilcoxon exact rank-sum test). Gray dashed lines indicate the artifact threshold of 0.1, which differs from that used in (G) and was determined based on the ventral-view measurements of PHMs^27^. Scale bars, (A, B) 50 μm, (C, E) 200 μm.

**Figure S4.**
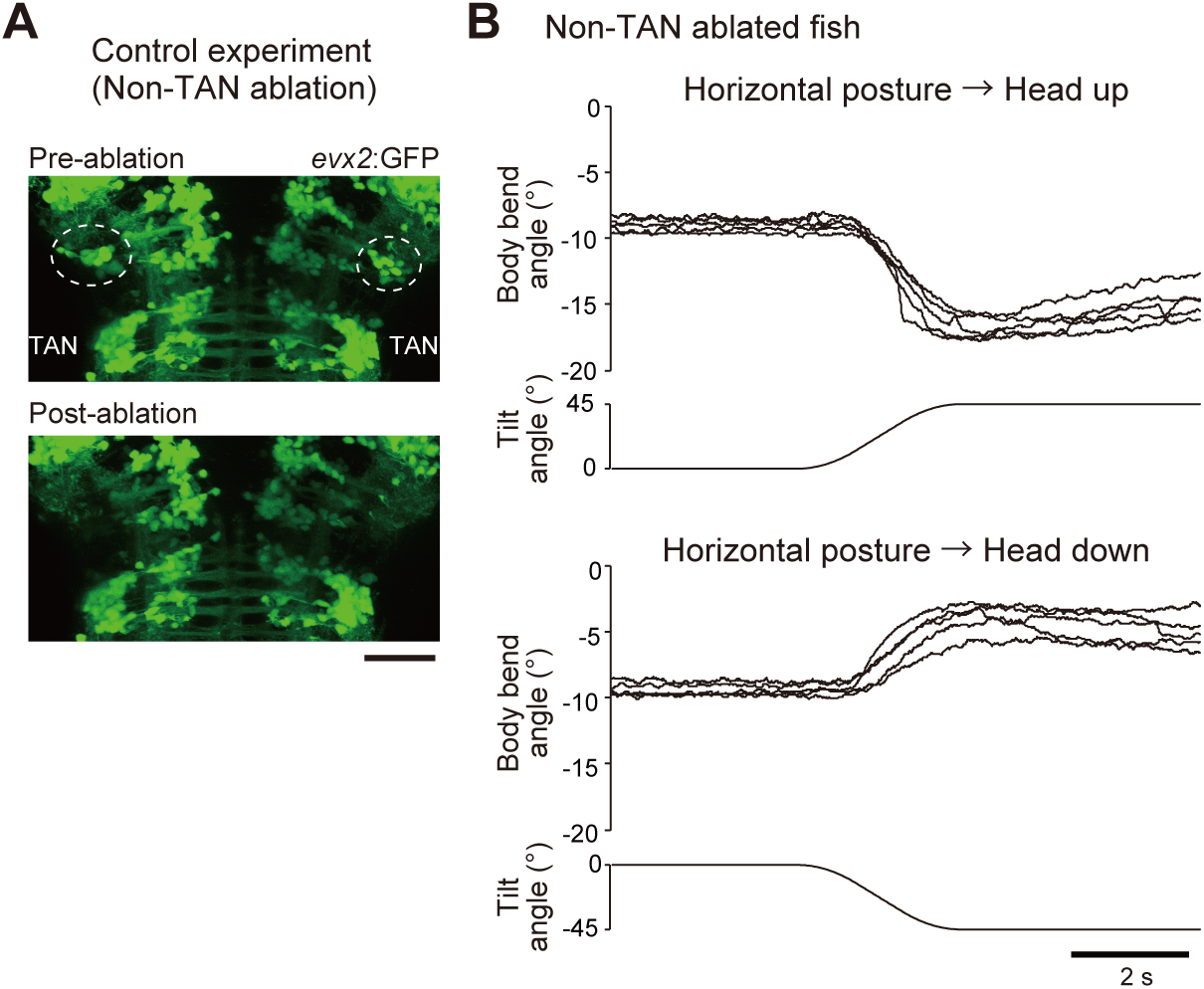
Body flexion in non-TAN-ablated animals. (A) Confocal stacked images of a Tg(*evx2*:GFP) fish before and after laser ablation of bilateral non-TAN neurons. Scale bar, 50 μm. (B) Body bend angle traces of a non-TAN-ablated fish in response to tilt stimuli under head-embedded conditions. Five trials for each tilt stimulus condition from the same fish are shown.

**Figure S5.**
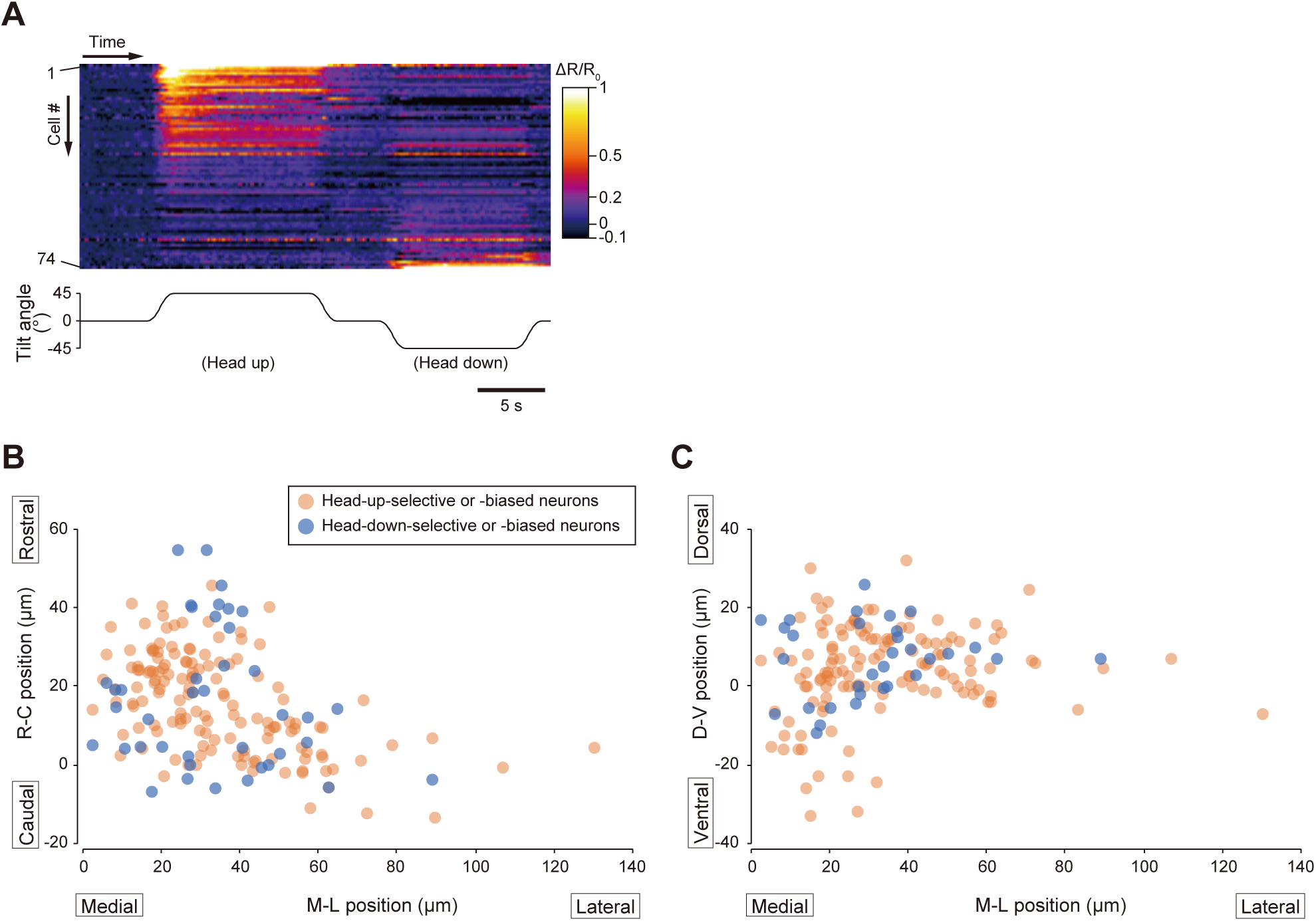
Characterization of Ca^2+^ responses of nMLF neurons. (A) Color-coded ΔR/R_0_ (changes in green/red ratio) time-series of 74 nMLF neurons from 5 fish in response to pitch tilts. Neurons are sorted based on the difference between head up/down tilt-induced ΔR/R_0_ amplitudes. (B, C) Spatial distribution of head-up-selective or -biased (orange; 143 cells) and head-down-selective or -biased (blue; 41 cells) neurons along the medio-lateral and rostro-caudal plane (B), and the medio-lateral and dorso-ventral plane (C). Positions were normalized relative to the midline (medio-lateral axis) and the position of MeLm (rostro-caudal and dorso-ventral axes).

**Figure S6.**
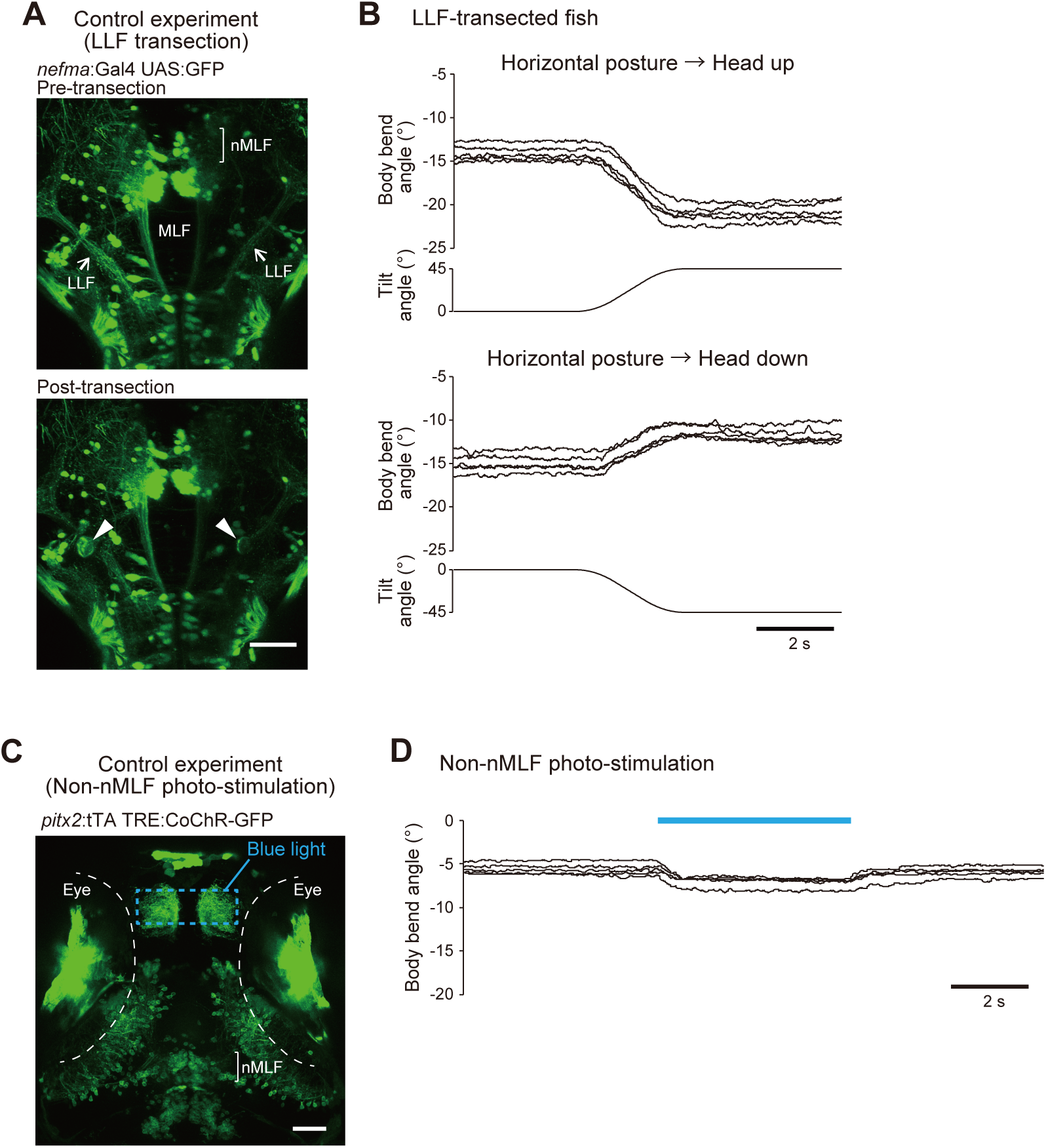
Axon transection and optogenetic activation of non-nMLF neurons. (A, B) Axon transection at lateral longitudinal fasciculus (LLF). (B) Confocal stacked images of a Tg(*nefma*:Gal4); Tg(UAS:GFP) fish before and after bilateral LLF transection at rhombomere 1. (C) Body bend angle traces of a LLF-transected fish in response to pitch tilts under head-embedded conditions. Five trials for each tilt stimulus condition from the same fish are shown. (C, D) Optogenetic activation of non-nMLF neurons. (D) Confocal stacked image of a Tg(*pitx2*:tTA); Tg(TRE:CoChR-GFP) fish. The blue dotted rectangle is the area illuminated with blue light. White dotted lines indicate the eye boundaries. (E) Body bend angle traces upon blue light illumination (blue bar, 5 s) under head-embedded conditions. Four trials in a single fish are shown. Scale bars, 50 μm.

**Movie 1. Ventral flexion in response to a head-up pitch tilt**

Behavior in response to a head-up tilt under head-embedded conditions. Raw (left) and rotated (right) movies are shown side by side. In the rotated movie, image registration was performed using the head region to facilitate visualization of the body shape. The movie is played at real speed.

**Movie 2. Dorsal flexion in response to a head-down pitch tilt**

Same as Movie 1, but behavior in response to a head-down tilt.

**Movie 3. Ventral flexion upon optogenetic activation of nMLF neurons**

Ventral flexion upon blue light illumination around the area of nMLF neurons in a Tg(*pitx2*:tTA); Tg(TRE:CoChR-GFP) fish under head-embedded conditions. The movie is played at real speed.

## Table

**Table S1.** List of reagents and transgenic lines.

| REAGENT or RESOURCE | SOURCE | IDENTIFIER |
| --- | --- | --- |
| **Antibodies** |  |  |
| S58 | DSHB | Cat# s58, RRID: AB_528377 |
| Goat anti-Mouse IgG (H+L) Highly Cross-Adsorbed Secondary Antibody, Alexa Fluor 488 | Thermo Fisher Scientific | Cat# A-11029, RRID: AB_2534088 |
| **Chemicals, peptides, and recombinant proteins** |  |  |
| Ethyl 3-aminobenzoate methanesulfonate salt (MS-222) | Sigma Aldrich | Cat# A5040 |
| Agarose-LM | Nacalai Tesque | Cat# 01161-12 |
| Cal-520 Dextran conjugate MW 10,000<br>Dextran, Tetramethylrhodamine, 3000 MW, Anionic, Lysine Fixable | AAT bioquest<br>Thermo Fisher Scientific | Cat# 20601<br>Cat# D3308 |
| **Experimental models: Organisms/strains** |  |  |
| Tg(*smyhc2*-hs:tdTomato-jGCaMP8s) | This paper |  |
| Tg(*smyhc2*-hs:loxP-RFP-loxP-DTA) | Sugioka et al., 2023<sup>27</sup> |  |
| Tg(*smyhc2*:loxP-RFP-loxP-DTA) | This paper |  |
| Tg(*tbx2b*-hs:Cre) | Sugioka et al., 2023<sup>27</sup> |  |
| Tg(*zic1*-hs:Cre) | This paper |  |
| Tg(*evx2*-hs:GFP) | Kawano et al., 2022<sup>64</sup> | ZFIN ID: ZDB-CRISPR-230523-1 |
| Tg(*nefma*-hs:Gal4);Tg(UAS:GFP) | Liu et al., 2020<sup>65</sup> | ZFIN ID: ZDB-ALT-200623-2 |
| Tg(*pitx2*-hs:tTA) | This paper |  |
| Tg(TRE:CoChR-GFP) | Uemura et al., 2020<sup>66</sup> |  |
| Tg(*pitx2*-hs:Dendra2) | Sugioka et al., 2023<sup>27</sup> |  |
| Tg(*actc1b*:GFP)<br>(previously known as Tg(α-actin:GFP) | Higashijima et al., 1997<sup>74</sup> | ZFIN ID: zf13Tg |

Note that “-hs” is omitted in the Results and Figures/Legends.

**Table S2.** List of key optomechanical components.

| REAGENT or RESOURCE | SOURCE | IDENTIFIER |
| --- | --- | --- |
| **Software and algorithms** |  |  |
| Kinesis | Thorlabs | https://www.thorlabs.com/newgrouppage9.cfm?objectgroup_id=10285 |
| FlyCapture2 | Teledyne FLIR | https://flycap2-viewer-release.software.informer.com |
| ImageJ/Fiji<br>python | NIH<br>Python Software<br>Foundation | https://imagej.net<br>https://www.python.org/ |
| R | R Core Team | https://www.r-project.org/ |
| Excel | Microsoft | https://www.microsoft.com/en-us/microsoft-365/excel |
| **Other** |  |  |
| Motorized rotation stage | Thorlabs | DDR100/M |
| Digital mirror device | Mightex | Polygon400 |
| Confocal scanner unit | Yokogawa | CSU-X1 |
| Image splitting optics | Hamamatsu Photonics | W-VIEW GEMINI, A12801-01 |
| Digital camera | Hamamatsu Photonics | ORCA-Flash4.0 V3, C13440-20CU |
| Light source | Excelitas Technologies | X-Cite exacte |
| Light source | Prizmatix | UHP-LED-470 |
| Digital camera | Teledyne FLIR | GS3-U3-23S6M-C |
| Digital camera | The Imaging Source | DMK33UX174 |
| Objective lens | Olympus/Evident | 10×/ NA 0.3, UPLANFLN |
| Objective lens | Olympus/Evident | 10×/ NA 0.3, UMPLANFLN |
| Objective lens | Olympus/Evident | 40×/ NA 0.8, LUMPLANFLN |
| Lens | Tamron | M118FM50 |
| close-up extension tube | Tamron | MR-set |

